# Mapping the functional differentiation and interactions among the inferior, medial frontal and posterior temporal cortex in semantic control

**DOI:** 10.1101/2025.04.28.650132

**Authors:** Jiaxuan Liu, Xiuyi Wang, Hanqing Zhang, Junjie Yang, Xiaowei Gao, Zhongqi Li, Yaling Wang, Zhe Hu, Junjing Li, Wanjing Li, Yien Huang, Jiali Chen, Lizhang Zeng, Xuchu Weng, Binke Yuan

**Affiliations:** Key Laboratory of Brain, Cognition and Education Sciences, Ministry of Education, China: Institute for Brain Research and Rehabilitation, South China Normal University, Guangzhou 51063, China; State Key Laboratory of Cognitive Science and Mental Health, Institute of Psychology, Chinese Academy of Sciences, Beijing 100101, China; Department of Psychology, University of Chinese Academy of Sciences, Beijing 100049, China; Philosophy and Social Science Laboratory of Reading and Development in Children and Adolescents (South China Normal University), Ministry of Education, Guangzhou 51063, China; Center for Studies of Psychological Application, South China Normal University, Guangzhou 51063, China; Guangdong Key Laboratory of Mental Health and Cognitive Science, South China Normal University, Guangzhou 51063, China

**Keywords:** semantic control, inferior frontal gyrus (IFG), posterior middle temporal gyrus (pMTG), dorsomedial prefrontal cortex (dmPFC), functional magnetic resonance imaging (fMRI), transcranial magnetic stimulation (TMS), dynamic causal model (DCM)

## Abstract

Semantic control refers to the ability to flexibly retrieve and manipulate stored knowledge to support context-appropriate behavior. A left-lateralized network comprising the left inferior frontal gyrus (IFG), posterior middle temporal gyrus (pMTG), and dorsal medial prefrontal cortex (dmPFC) has been consistently implicated in this process. While previous studies have established the necessity of the IFG and pMTG in semantic control, the causal role of the left dmPFC remains unclear. Additionally, it is unknown whether each of these three regions exhibits internal functional differentiation and how they interact to support semantic control. To address these questions, we combined task-based functional magnetic resonance imaging (fMRI) with fMRI-guided transcranial magnetic stimulation (TMS). We found that dmPFC, like IFG and pMTG, is causally involved in semantic control. All three regions exhibited a consistent anterior–posterior functional gradient: anterior subregions were selectively engaged during high-demand semantic processing, whereas posterior subregions responded to both easy and hard tasks. Furthermore, combined activation patterns of these regions better predicted the behavioral differences between hard and easy semantic tasks compared to the activation patterns of any single region. Semantic control modulated both the autoinhibition within individual regions and the functional connectivity among them, suggesting these regions operate in a coordinated network rather than in isolation. These findings advance our understanding of the neural architecture supporting flexible semantic behavior.

**Significance Statement:** Understanding how the brain supports flexible semantic behavior is critical for both cognitive neuroscience and clinical neuropsychology. While prior studies have consistently implicated the left IFG, pMTG, and dmPFC in semantic control, the causal contribution of the dmPFC and the functional dynamics among these regions have remained unclear. This study provides the first causal evidence for the dmPFC’s role in semantic control, reveals functional differentiation within the IFG and pMTG, and shows that these regions interact as an integrated network. These findings challenge the notion of functionally homogeneous control nodes and highlight a topographically organized, interactive system underlying controlled semantic retrieval. This work refines our mechanistic understanding of semantic control and may inform clinical models of language and conceptual deficits.

## Introduction

Semantic control is the capacity to flexibly retrieve and manipulate stored knowledge, especially when automatic processing is insufficient for demanding semantic tasks (Jefferies, 2013; Lambon Ralph et al., 2017). It supports context-appropriate meaning selection during reading comprehension (Whitney et al., 2011a; Hoffman & Tamm, 2020; Gao et al., 2022; Liang et al., 2024), facilitates the retrieval of weak associations and the suppression of dominant ones during communication (Whitney et al., 2011b, 2012; Hallam et al., 2016; X. Wang et al., 2024a), enables flexible conceptual inference through feature- and rule-based reasoning (Stawarczyk & D’Argembeau, 2015; Davey et al., 2016; Chiou et al., 2018, 2023), and contributes to emotional regulation via cognitive reappraisal (Dörfel et al., 2014). Large-scale meta-analyses have consistently identified a predominantly left-lateralized semantic control network (SCN), comprising the left inferior frontal gyrus (IFG), posterior middle temporal gyrus (pMTG), and dorsomedial prefrontal cortex (dmPFC), which are reliably recruited under high semantic control demands (Noonan et al., 2013; Lambon Ralph et al., 2017; Jackson, 2021). However, the causal roles and functional differentiation for each region and inter-regional interactions remains unclear.

Convergent causal evidence already demonstrates the necessity of IFG and pMTG, as inhibitory non-invasive brain stimulation (NIBS) targeting either region selectively impairs performance on high control demand semantic tasks, while sparing easier semantic or non-semantic tasks (Whitney et al., 2011b, 2012; Davey et al., 2015; Hallam et al., 2016; Alonso et al., 2024). In contrast, the dmPFC’s causal role remains untested. Nevertheless, multiple lines of evidence implicate its involvement in semantic control. First, within the medial prefrontal cortex gradient, the dmPFC is situated between anterior cingulate cortex (ACC) supporting domain-general control and ventromedial sectors involved in default mode integration, suggesting a topographic nexus for semantic control (Chiou et al., 2023; X. Wang et al., 2024a). Second, its cross-domain activation is increasingly interpreted as goal-directed selection of conceptual knowledge rather than social cognition per se (Binder & Desai, 2011; Binney & Ramsey, 2020; Diveica et al., 2021; Hodgson et al., 2021). Supporting this view, recent evidence shows that the dmPFC constructs compressed, task-relevant representations by filtering incoming social information (Wittmann et al., 2025), suggesting a computational mechanism analogous to semantic control. Crucially, the dmPFC shows strong functional and structural connections with both semantic representation and control regions. The ventral anterior temporal lobe (vATL)—an established semantic representation hub—shows strong functional connectivity with IFG, pMTG and dmPFC (Jackson et al., 2016), while the inferior longitudinal fasciculus (ILF), inferior fronto-occipital fasciculus (IFOF) and frontal aslant tract (FAT) furnish direct white-matter links among these regions (Duffau, 2015; Nugiel et al., 2016; Rojkova et al., 2016; Pascual-Diaz et al., 2020; Burkhardt et al., 2022). Disruption of the IFOF and FAT, as shown in intraoperative stimulation studies to elicit lexico-semantic errors (Corrivetti et al., 2019). Moreover, semantic-aphasia patients commonly exhibit structural and functional disconnections implicating the dmPFC (Souter et al., 2022). These findings collectively support the view that the dmPFC is an integral SCN node, motivating a direct causal test of its necessity with the prediction that inhibitory stimulation will selectively impair performance under high semantic control demands.

A central issue is whether the three putative SCN regions operate as functionally homogeneous units or exhibit systematic internal heterogeneity. Research has portrayed these regions as interchangeable substrates dedicated exclusively to semantic control (Noonan et al., 2013; Hallam et al., 2016; Jackson, 2021). More recent work, however, links cortical topography to functional specialization and increasingly supports a gradient-based account: domain-general control zones lie posteriorly, near unimodal sensorimotor cortex, whereas semantic-specific control zones shift anteriorly toward the heteromodal apex of the cortical hierarchy (X. Wang et al., 2024a, 2024b). This gradient is well documented in the IFG, where anterior-ventral subregions couple with semantic-specific networks and are recruited during controlled retrieval, whereas posterior-dorsal subregions align with multiple demand networks (MDN) to resolve competition and support domain general processing (Badre, 2005; Badre & Wagner, 2007; J. Wang et al., 2020; Fedorenko & Blank, 2020; Diveica et al., 2023). A similar pattern is observed in the pMTG and dmPFC, where demanding semantic association judgments preferentially activate anterior segments, whereas feature-matching and non-semantic control tasks recruit more posterior segments (Chiou et al., 2023; X. Wang et al., 2024a). Thus far, these gradients have been supported almost exclusively by correlational neuroimaging and connectivity studies, and no study has provided causal evidence that perturbing anterior versus posterior sub-regions produces distinct behavioral effects.

Another primary yet unresolved question is how these SCN regions interact to facilitate semantic control. A study combining repetitive transcranial magnetic stimulation (rTMS) and fMRI found that inhibition of the left IFG increases the left pMTG and pre-SMA activity under higher semantic control demands (Hallam et al., 2016). An fMRI-based dynamic causal modeling (DCM) has demonstrated that increased context integration demands enhance inhibitory influence from the anterior IFG to the posterior superior temporal sulcus/middle temporal gyrus (pSTS/pMTG; Hartwigsen et al., 2017). While these findings suggest dynamic interplay within the SCN, the effective connectivity among its three core nodes—particularly involving the dmPFC—has yet to be systematically examined.

To address these gaps, the current study employed a 2 × 2 factorial experimental design involving semantic and visual tasks under high and low demand conditions to precisely localize semantic control regions. To investigate their causal roles, we applied a virtual lesion approach using TMS to determine whether inhibiting each region would disrupt subjects’ behavioral performance in controlled semantic retrieval. The individualized TMS inhibition further enabled us to assess the functional distinctions within each region. Furthermore, to examine whether these regions function cooperatively or independently, we conducted machine-learning-based multivariate prediction analyses to assess whether combined activation patterns of these regions could better predict behavioral performance than activation patterns from a single region. If the regions cooperate, we further explore how they interact to facilitate controlled semantic retrieval. To this end, we implemented dynamic causal modeling (DCM) to investigate the excitatory and inhibitory regulations among them and the behavioral relevance.

## 2 Material and methods

### 2.1 Participants

For the fMRI experiment, thirty-five right-handed young adults were recruited (18 females; 20.86 ± 2.28 years; age range, 18–29). Among these participants, twenty-two (11 females; 21.27 ± 2.47 years; age range, 18–29) also took part in the subsequent TMS experiments. Ethical approval for both fMRI and TMS experiments was granted by the ethic committee of the Institute for Brain Research and Rehabilitation at South China Normal University (grant number: SCUU-BRR-2022-015). Before participating, all individuals provided written informed consent, which included detailed information about the experimental procedure, potential risks and side effects of TMS and fMRI, the benefits of the study, and assurances of confidentiality. Participants were compensated for their time, receiving 100 RMB per hour for the MRI scanning and 50 RMB per session for the TMS experiment.

### 2.2 Experimental design

The experimental framework comprised six sessions, as illustrated in Figure 1a. The first session was a TMS sensitivity test designed to screen out individuals with excessive pain sensitivity. This procedure ensured participants’ suitability for subsequent TMS experiments. During this session, participants first underwent a 40-second application of continuous Theta Burst Stimulation (cTBS) targeting the left IFG, followed by behavioral tasks to familiarize them with the experimental procedures. In the second session, participants underwent MRI scans, and the task activation results were used to select precise stimulation sites for subsequent TMS experiments (see the Methods section for details). From the third to the sixth sessions, participants who passed the TMS sensitivity test received cTBS on one of four brain regions: the left IFG, pMTG, dmPFC, or the Vertex. The vertex (i.e., Cz point defined in the international 10-20 system) served as an active control site (Koessler et al., 2009; Rojas et al., 2018; Q. Zhang et al., 2019). To ensure an accurate assessment of TMS effects, the stimulation order was counterbalanced across participants. After each stimulation, participants took a short 5-minute break before completing the behavioral tasks. Additionally, a minimum five-day interval between TMS sessions was implemented to allow for the dissipation of any residual effects from previous sessions, further enhancing the reliability of the results.

**Figure 1.**
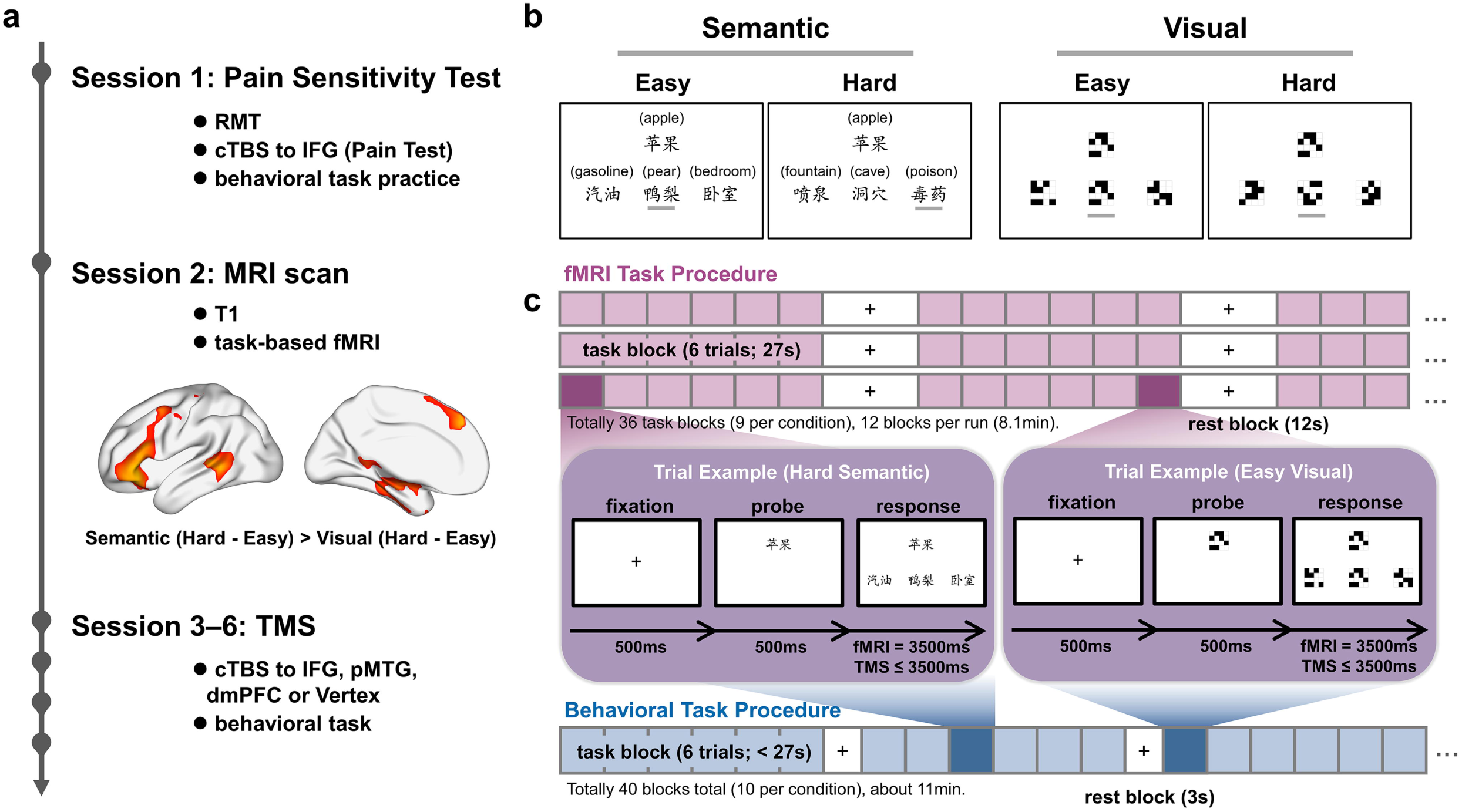
Experimental sessions and paradigm. **(a)** The experiment comprised six sessions: a pain sensitivity test (Session 1), an MRI scan (Session 2), and four TMS sessions (Sessions 3–6). Semantic control regions were functionally localized using the contrast (Semantic Hard - Easy) > (Visual Hard - Easy) from the fMRI task (see Figure 3 for activation details). The three regions with the strongest activation—the left IFG, dmPFC, and pMTG—were selected as TMS targets. In the TMS sessions, cTBS was applied to each target region or to the vertex (control site), followed by behavioral testing. **(b)** Task stimuli include semantic and visual conditions with easy (low control demand) and hard (high control demands) trials. In the semantic task, probe-target word pairs were either strongly associated (easy: e.g., *apple-pear*) or weakly associated (hard: e.g., *apple-poison*). In the visual task, probe-target pairs were either identical (easy) or rotated 180 degrees (hard) from the probe. **(c)** Task procedure in the fMRI session and TMS sessions. The fMRI session (pink) included 36 task blocks (9 per condition across three runs), each with six trials. Each TMS session (blue) comprised 40 blocks (10 per condition), also with six trials per block. IFG, inferior frontal gyrus; pMTG, posterior middle temporal gyrus; dmPFC, dorsomedial prefrontal cortex; cTBS, continuous theta burst stimulation.

### 2.3 Task and stimuli

To identify brain regions involved in semantic control, we utilized a semantic judgment task and a visual judgment task. The visual judgment task served as a control condition. Both tasks were adapted from previous fMRI and TMS studies (Whitney et al., 2011b; Q. Zhang et al., 2019; Chiou et al., 2023). We employed a 2 × 2 factorial design, orthogonally manipulating the type of judgment task (semantic vs. visual) and difficulty (easy vs. hard). This design yielded four distinct task conditions: easy semantic, hard semantic, easy visual, and hard visual condition.

As illustrated in Figure 1b, each trial comprised four items: a probe item, the target item, and two unrelated distractor items. In the semantic judgement task, participants were required to select the target word that was most closely related in meaning to the probe word from a set of three options. All stimuli consisted of Chinese concrete noun words. To minimize repetition, the same probe word was not repeated within a session, and the position of the target words varied randomly across trials, with equal probability of appearing in any of the three positions. Task difficulty, i.e., cognitive control demand, was manipulated by varying the strength of association between the probe word and the target word. Weak semantic associations required higher levels of semantic control, making these conditions more challenging, whereas strong associations required less semantic control, rendering those conditions easier. The association strength of both probe-target and probe-distractor word pairs was quantified using the Baroni-Urbani association measure derived from sentence windows in the Chinese Lexical Association Database (Lin et al., 2019), with distractor words having an association strength of zero with the probe. For all six sets of task stimuli (including one used in fMRI and five used in TMS tasks), the mean association strength of probe-target word pairs differed significantly between hard and easy condition in each set (independent t-tests: *ps* < 0.001; detailed in Table S2). A two-way ANOVA revealed no significant differences in stroke (Sun et al., 2018), lexical frequency (Cai & Brysbaert, 2010), familiarity, and imageability of the words across high and low semantic control conditions and sessions (see Table S1 and S3). Familiarity and imageability scores for the words were rated by an independent group of 20 participants using a 7-point scale. In the visual judgment task, participants were required to select a target pattern from three options that matched a provided probe pattern. The stimuli consisted of seven black squares placed randomly within a 4 × 4 grid of white squares. In the easy condition (low control demand), the target pattern was identical in both shape and orientation to the probe pattern. In contrast, in the hard condition (high control demand), the target shared the same shape as the probe but was rotated by 180 degrees. The position of the target patterns varied randomly in each trial, with each of the three positions equally probable.

### 2.4 Task procedure

#### 2.4.1 fMRI task in scan session

Participants completed a mini-block design task in the fMRI scanner, consisting of three runs. Each run included three task blocks per condition, totaling 12 blocks across all conditions. Each task block contained six trials of the same condition, resulting in a total of 216 trials across the entire task. Each trial began with a 0.5 second fixation, followed by the presentation of the probe item for 0.5 second. Subsequently, all items were displayed with a fixed duration of 3.5 seconds. Task blocks were presented in a random order, with 12 seconds fixation blocks interspersed between them. Additionally, each run began and ended with separate 15-second fixation periods, avoiding signal instability or loss. The total duration of a single run was 8.1 minutes.

Participants were required to select the target items by pressing one of three designated buttons on a magnetic resonance-compatible response box. Stimuli were presented using MATLAB (www.mathworks.com) and the Psychtoolbox-3 distribution (www.psychtoolbox.org), displayed on an MRI-specialized projection screen (1920 × 1080 resolution). Participants viewed the stimuli on the screen through a mirror mounted on the radio frequency coil.

#### 2.4.2 Behavioral task in TMS sessions

Participants who passed the TMS sensitivity test completed four separate experimental sessions on different days, with the order of behavioral tasks randomized. Each session included 10 task blocks per condition, totaling 40 task blocks. Each task block comprised six trials of the same condition, resulting in a total of 240 trials per task. Each trial began with 0.5-second fixation, followed by the presentation of the probe item for 0.5 second, and then all items were displayed with a maximum of 3.5 seconds. Task blocks were presented in a random order, with 3-second fixation blocks interspersed between them. The behavioral tasks were self-paced and lasted approximately 11 minutes. The mean response times for each stimulation site were as follows: IFG, 675.16 ± 50.33 ms; pMTG, 672.56 ± 52.37 ms; dmPFC, 671.51 ± 49.54 ms; Vertex, 656.98 ± 49.56ms). Stimuli were presented using MATLAB (www.mathworks.com) and the Psychtoolbox-3 distribution (www.psychtoolbox.org). Participants sat in front of a computer screen and select the target items by pressing one of the numeric keys (1, 2, or 3).

### 2.5 MRI data acquisition

MRI data were collected using a Siemens 3-Tesla Prisma Fit scanner equipped with a 64-channel head coil (Siemens, Erlangen, Germany), at the Brain Imaging Center of South China Normal University. Except for one participant whose resting-state fMRI data was not scanned, all participants completed T1 imaging scans, task-state fMRI scans, and resting-state fMRI scans. T1-weighted anatomical images were acquired using a 3D Magnetization Prepared Rapid Gradient Echo (MPRAGE) sequence [repetition time (TR) = 1900 ms; echo time (TE) = 2.27 ms; inversion time (TI) = 1100 ms; bandwidth = 190 Hz/Px; flip angle = 7°; field of view (FOV) = 256 mm; matrix size = 256 × 256 × 208; voxel size = 1 × 1 × 1 mm^3^ in-plane acceleration factor (iPAT) = 2]. Functional MRI data were obtained with an echo-planar imaging (EPI) pulse sequence [TR = 1500 ms; TE = 30 ms; flip angle = 90°; FOV = 192 mm; voxel size = 3 × 3 × 3 mm^3^; 43 slices; bandwidth = 2520 Hz/Px; in-plane acceleration factor (iPAT) = 2]. For the current study, a series of 324 functional volumes were acquired for task-based fMRI, and 240 functional volumes were acquired for resting-state fMRI.

### 2.6 MRI data analysis

#### 2.6.1 preprocessing

Task-based fMRI data preprocessing was performed using SPM12 software (Statistical Parametric Mapping; www.fil.ion.ucl.ac.uk/spm/software/ spm12/). The T1 images were segmented into gray matter (GM), white matter (WM), and cerebrospinal fluid (CSF) partitions. The functional images were preprocessed with the following steps: 1) Removement of the first 10 volumes and last 10 volumes; 2) Correction of slice timing; 3) Head motion correction; 4) Co-registration of each T1 image to a functional image; 5) Normalization to Montreal Neurological Institute (MNI) space with solution of 3 × 3 × 3 mm; 6) Spatial smoothing with an isotropic Gaussian kernel (FWHM = 8 mm). No participant exhibited head motion > 3 mm maximum translation or 3° maximum rotation.

#### 2.6.2 Whole-brain general linear model (GLM) analysis

To identify brain regions responsible for semantic control, a whole-brain general linear model (GLM) analysis was conducted using SPM12 (7771). To enhance statistical power, data from the three runs were concatenated prior to the GLM, with n - 1 additional (unconvolved) regressors included to model run-specific means. For each participant, an experimental design matrix was created, specifying the onset times and durations of each task block as regressors. This matrix was convolved with a canonical hemodynamic response function to model the BOLD response to task stimuli. Additionally, the six head motion parameters were incorporated into the design matrix to mitigate movement-related artifacts. A high-pass filter was applied to eliminate low-frequency noise.

The analysis focused on the interaction contrast Semantic (Hard - Easy) > Visual (Hard - Easy), which isolates brain regions specifically engaged in high-level semantic control processing. This contrast was entered into a random-effect model, and a one-sample t-test was performed on the summary statistic for multi-subject analysis. The statistical threshold was set at *p* < 0.01, corrected for multiple comparisons using the False Discovery Rate (FDR) method at the voxel level, with a minimum cluster size of 50 voxels required for significance. To ensure the relevance of the findings, a mask of semantic positive activation (hard semantic condition versus rest baseline, *p* < 0.01, FDR corrected at the voxel level, cluster size = 50 voxels) was applied to the results of the primary contrast.

In the results, the three most significant group-level clusters — the left IFG, pMTG, and dmPFC — were identified and selected for subsequent analyses. These regions served as targets for cTBS targets and were further examined using DCM analysis. Detailed methods for these analyses are described in their respective sections.

#### 2.6.3 Multivariate machine-learning-based prediction analysis for behavioral performance based on activation patterns

To investigate whether core semantic control regions function collaboratively or independently, we constructed multivariate machine learning models to test whether combined activation patterns from three regions could better predict participants’ semantic task performance compared to patterns from single regions. Specifically, if the combined model significantly outperformed single-region models, it would suggest collaborative functionality among these regions; otherwise, the regions might function independently.

We employed the relevance vector regression (RVR) method, based on the sparse Bayesian learning framework with a linear kernel function (Tipping, 2001). Automatic relevance determination (ARD) was applied to impose independent Gaussian priors on the weights of basis functions associated with each training sample, enabling the model to identify the most relevant samples for prediction (i.e., *relevance vectors*). Hyperparameters controlling the prior variance of each sample’s weight were optimized via the type-II maximum likelihood method (Berger, 1985). This procedure generates sparse models by retaining only critical samples, enhancing generalization performance while minimizing model complexity (Tipping, 2001). Prior studies have demonstrated the efficacy of RVR methods in predicting brain-behavioral/cognitive mapping (Cui & Gong, 2018; Yuan et al., 2019, 2023).

Four models were constructed: a combined model integrating activation patterns from all three regions, and three single-region models (IFG, dmPFC, and pMTG). The feature inputs for these models consisted of voxel activation beta values derived from the semantic hard > easy contrast, with all manually specified parameters held constant. The predicted label was the difference in RTs between the hard and easy semantic condition (ΔRT). Leave-one-out cross-validation (LOOCV) was employed to assess the generalizability of these models. In each LOOCV iteration, one participant was left out as the test set, while the remaining data served as the training set for model prediction.

To address the challenge of high-dimensional voxel-based features relative to the limited sample size, a univariate feature-filtering approach based on Pearson correlation was implemented (Guyon & Elisseeff, 2003; Di Martino et al., 2008; Pereira et al., 2009; Dosenbach et al., 2010). For each LOOCV iteration, feature selection and parameter optimization were performed exclusively within the training set. Voxels were ranked in ascending order of their p-values derived from the Pearson correlation between their beta values and ΔRTs. To determine the optimal number of voxels for inclusion, a grid search was conducted based on p-value thresholds ranging from 0.01 to 1 in increments of 0.01. The optimal threshold was selected based on the highest prediction performance, i.e., the largest Pearson correlation coefficient between predicted and actual labels across all thresholds tested.

To comprehensively evaluate model generalization performance, we employed three different LOOCV-based metrics. First, we computed the Pearson correlation coefficient (r) between predicted and actual ΔRT values, assessing significance through a permutation test with 1000 iterations. For each permutation, predicted labels were randomized, and the RVR process was repeated to generate a null distribution of Pearson r values, determining the empirical significance of the observed correlation. Second, prediction errors, including absolute error (AE) and squared error (SE), were computed across all test samples within the LOOCV framework. Differences in prediction errors between models were statistically compared using paired t-tests, with statistical significance through permutation testing (5,000 iterations) and Bonferroni correction for multiple comparisons. Finally, To account for differences in model complexity and overfitting risks, a Bayesian model selection (BMS) approach was employed (Myung & Pitt, 2018). In each LOOCV iteration, the *negative log marginal likelihood* of each model was computed using the *Laplace approximation* method within the RVR framework (Tipping, 2001). By averaging these values across all iterations, an overall measure of model evidence was obtained. The log Bayes Factor (logBF) was then calculated to assess the relative performance of different models in balancing prediction accuracy and complexity, providing a robust measure of overall model performance.

#### 2.6.4 Effective connectivity analysis: DCM analysis

To investigate the communication mechanisms among the three core semantic control regions, we employed Dynamic Causal Modeling (DCM) with Parametric Empirical Bayes (PEB) framework (Zeidman et al., 2019a, 2019b). Unlike conventional DCM, which requires separate second-level analysis, the PEB framework enables hierarchical Bayesian modeling at the group level, improving statistical efficiency and allowing for direct estimation of individual variability. We hypothesized that higher semantic control demands would reshape the interaction patterns among these regions by modulating their effective connectivity. To account for individual differences in activation, we identified the closest individual-level peaks within an 8-mm radius spherical region centered around the group-level peaks. Time series were extracted form supra-threshold voxels as the first eigenvariate within a sphere of 6 mm radius around the individual-level peaks in the semantic contrast (hard > easy, at a threshold of uncorrected *p* < 0.05). Participants with noisy or missing data in one or two regions were not excluded; instead, the threshold was gradually lowered to p < 0.30 (uncorrected) to ensure the extraction of time series data. This approach leverages the Bayesian hierarchical modeling framework, which appropriately down-weights the contribution of participants with noisy data to the group effect, ensuring that even participants with weal response in one brain region contribute useful information about other regions and intersubject variability (Zeidman, et al., 2019a).

##### Model specification and parameter estimation on individual level

For each participant, a full DCM model was specified, with connectivity parameters described by three matrices (A, B, and C). The A matrix quantified the intrinsic effective connectivity among the three regions, including reciprocal connections between regions and self-connections within each region. Since experimental inputs were mean-centered, the values in matrix A represent the average effective connectivity across semantic conditions. The matrix B captured how the effective connectivity, as defined in matrix A, was modulated by experimental conditions of interest. To thoroughly investigate interaction patterns among the three brain regions, self-connections and bidirectional interregional connections were all specified in matrix B. The matrix C represented the sensitivity of each region to driving inputs. In our model, all semantic trials were considered driving inputs exerting for all three regions, while the hard semantic trials alone served as the modulatory input.

The full DCM model for each participant was estimated using the Variational Laplace approach (K. Friston et al., 2007; Zeidman et al., 2019a). As a diagnostic measure, the explained variance for each DCM model we calculated, with an average of 10.84% across participants. This relatively low explained variance was expected, as the model focused solely on semantic task-related signals, which constitute only a portion of the entire time series. By excluding control conditions and rest periods, a significant portion of the data was treated as noise, limiting the model’s explanatory power. Consequently, all participants were included in the group-level analysis to ensure a comprehensive representation of between-subject variability.

##### Group-level analysis based on parametrical empirical Bayes (PEB) framework

The DCM-PEB framework was used to assess the group effect on effective connectivity, integrating a hierarchical model with random effects on connection parameters (K. J. Friston et al., 2016). This approach accounts for both the expected values and the covariances of the parameters, ensuring that subjects with noisy data and uncertain parameter estimates are appropriately down-weighted. Thus, more precise parameter estimates exert greater influence on the group-level results (Zeidman et al., 2019b).

The covariance design matrix included the following components: (1) a “mean” regressor, representing the group mean effect as a column of ones for all participants; (2) a “residual variance (RV)” regressor, capturing individual variability in semantic control demand during the hard semantic condition by regressing the mean inverse efficiency score (i.e., reaction time divided by accuracy) under hard semantic condition (IE_SH_) on the mean inverse efficiency score under easy semantic condition (IE_SE_). This approach integrates both speed and accuracy, providing a nuanced performance metric while controlling general semantic ability; (3) covariates of no interest, specifically gender and age. Separate PEB analyses were conducted for matrices A, B, and C to maintain focus and clarity, with particular emphasis on the group mean effect and individual variability in semantic control demand.

Following the estimation of group-level parameters for the full DCM model, Bayesian model reduction (BMR) was applied to compare the free energy of the full model with that of reduced models. Reduced models were generated by systematically deactivating one or more parameters through an automated greedy search. Bayesian model average (BMA) was then performed, involving the calculation of a weighted average of parameters across 256 models obtained from the BMR (K. Friston & Penny, 2011; Rosa et al., 2012; K. J. Friston et al., 2016). We focused on parameters in the BMA model with strong evidence for their presence, defined as a posterior probability exceeding 0.95 for a parameter being present rather than absent (Zeidman, et al., 2019b).

### 2.7 TMS protocol

The continuous theta-burst stimulation (cTBS) protocol consists of a 40-second train of uninterrupted TBS, delivering three pulses at 50 Hz repeated every 200 ms, totaling 600 pulses (Huang et al., 2005). We chose cTBS due to its brief stimulation period and its ability to induce post-stimulus effects lasting approximately 50 minutes (Wischnewski & Schutter, 2015). The stimulation was delivered using a BrainSight neuro-navigation system (Rogue Research, Montreal, Canada) and a Magstim Rapid2 Stimulator with a 70 mm figure-eight coil (The Magstim Company, Whitland Dyfed, UK). The stimulation intensity was set at 80% of the individual’s resting motor threshold. RMT was defined as the lowest stimulation intensity required to elicit Motor Evoked Potentials (MEPs) exceeding 50 μV in the relaxed first dorsal interosseous muscle from at least 5 of 10 consecutive single-pulse stimulations of the right M1 area.The mean RMT across participants was 55.59 ± 4.81%. The mean stimulation intensities for the target regions were as follows: 44.14 ± 3.56% for the left IFG, 44.68 ± 3.94% for the left pMTG, 44.59 ± 3.75 % for the left dmPFC, and 44.86 ± 3.76 % for the vertex. This intensity level is considered safe according to recent guidelines (Rossi et al., 2021).

The target brain regions—left IFG, pMTG, dmPFC—were defined based on the results of the GLM activation analysis. To account for structural and functional variability among participants, the MNI coordinates of the stimulation sites were precisely adjusted to the individual-level peaks closest to the group-level peaks, ensuring they remained within the anatomical boundaries of the target regions as defined by the AAL atlas. In cases where individual-level peaks could not be identified within the target regions, the group-level peaks were used as substitutes. Electrical field simulations of the stimulation sites were performed using SimNIBS v.3.2.6 (Thielscher et al., 2015; Saturnino et al., 2018). Coil placement was optimized using the fast computational auxiliary dipole method (ADM; Gomez et al., 2021). The method was designed to quickly find a coil placement that maximizes the E-field strength of the stimulation sites. Additionally, practical considerations such as the participant’s pain tolerance and potential obstructions from the ears were taken into account when determining the final stimulation sites. The individual stimulation sites were mapped onto the standard ICBM152 template using BrainnetViewer (Xia et al., 2013), are illustrated in Figure 2. The average MNI coordinates for the target regions are as follows: IFG at [-49.77 ± 3.66, 30.41 ± 5.01, 6.14 ± 8.10], pMTG at [-58.55 ± 4.72, -46.91 ± 5.97, -0.68 ± 5.07], and dmPFC at [-4.50 ± 1.79, 37.23 ± 8.96, 42.73 ± 6.10]. The approximate location of the vertex is [-2, -15, 75].

**Figure 2.**
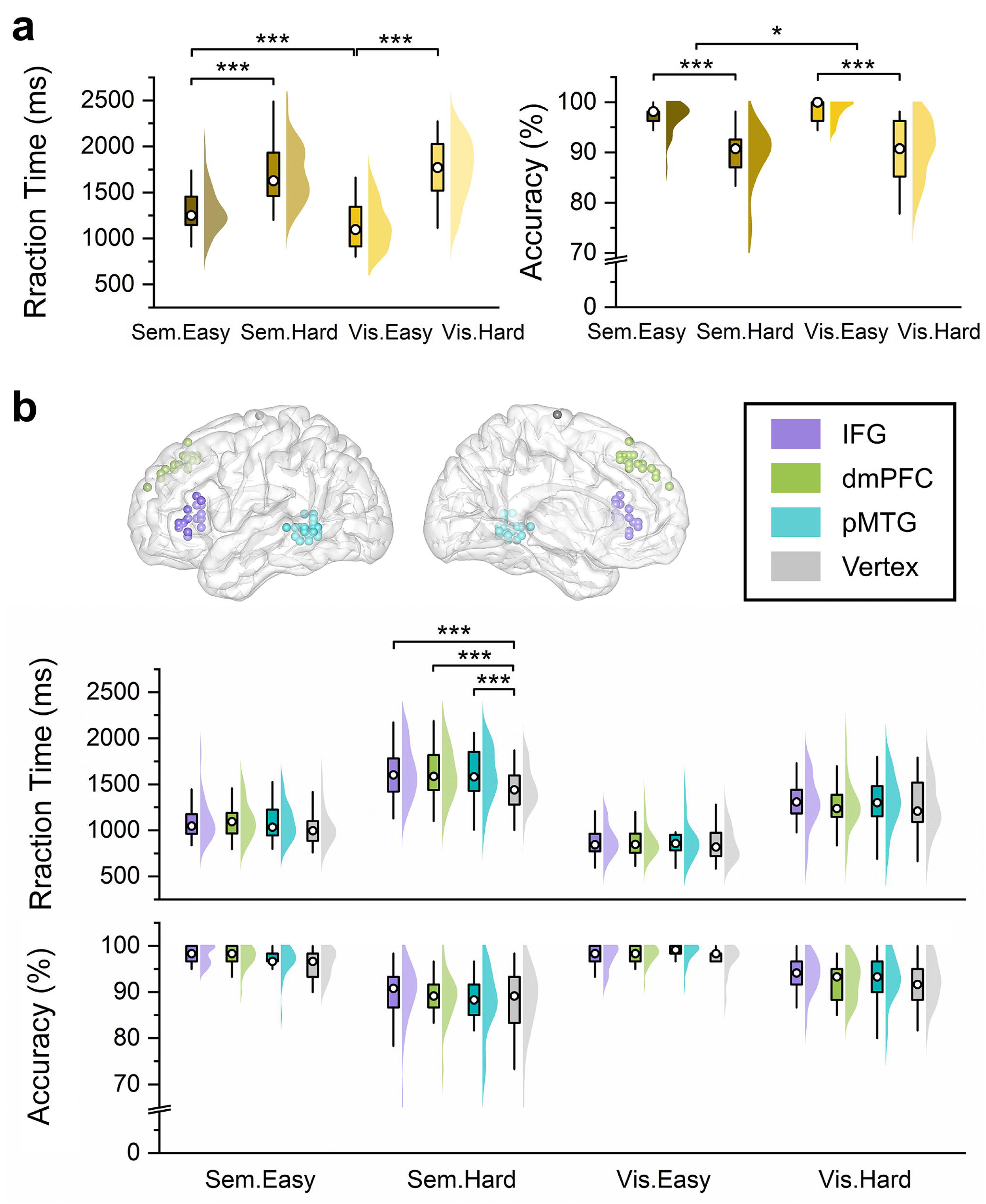
Behavioral results. **(a)** The behavioral results from the fMRI task are illustrated across four conditions, represented with a gradient from dark yellow to light yellow. The x-axis lists the conditions as Semantic Easy (Sem.Easy), Semantic Hard (Sem.Hard), Visual Easy (Vis.Easy), and Visual Hard (Vis.Hard). The y-axis represents behavioral measures, specifically mean RT (in milliseconds) and mean ACC (as a percentage). Data are visualized using Boxplots, with hollow circles indicating the medians and whiskers extending to 1.5 times the interquartile range. Half-violin plots are overlaid to show the distribution of the data, with color gradient from dark yellow to light yellow. **(b)** Locations of TMS stimulation are visualized on standard ICBM152 template surface. Each site for each subject is represented by a color-coded dot: the left IFG in purple, the dmPFC in green, the pMTG in teal, and the Vertex in grey. The behavioral results from the TMS session are depicted below. The x-axis displays the four conditions: Sem.Easy, Sem.Hard, Vis.Easy, and Vis.Hard, while the y-axis details mean RT and mean ACC. Boxplots show the median (denoted by hollow circles) and 1.5 times the interquartile range, with half-violin plots demonstrating the data distribution. Each region is represented by a specific color matching those on the standard brain surface. IFG, inferior frontal gyrus; pMTG, posterior middle temporal gyrus; dmPFC, dorsomedial prefrontal cortex; RT, reaction time; ACC, accuracy.

### 2.8 Statistical analysis of Behavioral data

Reaction times (RTs) and accuracy (ACC) were extracted for each participant and trial. Trials with incorrect response and outliers were excluded from the RT analysis. Specifically, trails with premature responses (< 200 ms), delayed responses (> 3500 ms), or those exceeding the mean by more than 3 SDs were removed. The mean RTs and ACC for each participant were then calculated for each condition. Subsequent statistical analysis at the group level was conducted using IBM SPSS Statistics (version 25.0.0.0).

To assess the differences in behavioral performance across task conditions in the fMRI task, we conducted a two-way repeated measures ANOVA, with task type (semantic vs. visual) and difficulty (hard vs. easy) as within-subject factors. Significant main effects or interaction were further analyzed using post-hoc tests to identify specific differences.

The impact of brain region stimulation on behavioral performance was investigated using three-way repeated measures ANOVA. The within-subject factors were TMS sites (IFG, pMTG, dmPFC, and control region vertex), task type (semantic vs. visual), and difficulty (easy vs. hard). If the assumption of sphericity was violated, as indicated by Mauchly’s test (p < 0.05), degrees of freedom were adjusted using the Greenhouse-Geisser correction. For any significant main effects or interaction, Bonferroni-corrected post hoc tests were performed to explore detailed analysis.

Given that portions of the left IFG, pMTG, and dmPFC were activated under both hard and easy semantic conditions in the fMRI task results (Figure 2a, b), participants were divided into two subgroups based on their stimulation sites within each region. This division aimed to investigate whether stimulation of different sub-regions lead to behavioral differences. Using the activation pattern from the easy semantic condition as a mask, we defined two groups per region. In the mid-IFG group (7 participants), stimulation sites were located within the mask, suggesting involvement in both easy and hard semantic condition. In contrast, the aIFG group (15 participants) had stimulation sites outside the mask, indicating potential engagement exclusively in hard semantic condition. Similarly, participants in the posterior pMTG group (12 participants) had stimulation sites within the mask, suggesting participating in both easy and hard semantic condition, whereas participants in the anterior pMTG group (10 participants) had stimulation sites outside the mask, likely involved only in hard semantic condition. The average MNI coordinates are: aIFG at [-49.29 ± 4.19, 32.57 ± 4.39, -3.00 ± 2.45], mid-IFG at [-50 ± 3.53, 29.40 ± 5.10, 10.40 ± 5.88], anterior pMTG at [-59.70 ± 5.74, -42.00 ± 4.90, -2.10 ± 3.18] and posterior pMTG at [-57.58 ± 3.65, -51 ± 2.86, 0.50 ± 6.11]. All stimulation sites for the dmPFC were located outside the mask, so no subgrouping was conducted for this region. For each subgroup, paired t-tests were performed to evaluate whether stimulation at the target site and the Vertex induced differences in RTs and ACC under each condition. To account for unequal group sizes and the overall small sample size, permutation testing (5,000 iterations) was employed to assess the statistical significance of the observed differences, with Benjamini-Hochberg FDR correction for multiple comparisons.

## 3 Results

In this study, we systematically investigated the roles and interactions of key brain regions in semantic control by manipulating task type and difficulty. Through a combination of fMRI and TMS experiments, we established the essential contributions of the left IFG, dmPFC, and pMTG to semantic control, confirming their causal involvement in these processes. To examine the functional differentiation within the left IFG and pMTG, as indicated by semantic activation patterns, we investigated whether TMS stimulation of specific subregions resulted in behavioral differences compared to Vertex stimulation across various task conditions. This analysis provided causal evidence for the distinct roles of the anterior and posterior subregions within these key areas, further refining our understanding of their functional organization. To determine whether these regions operate collectively or independently, we employed and compared multivariate machine learning models to predict behavioral differences based on the activation patterns of these three regions. DCM analyses further revealed that semantic difficulty modulates the interactions among these regions, highlighting their coordinated efforts in semantic control.

### 3.1 Behavioral performance of task-based fMRI

To evaluate whether our task design effectively manipulated the control demands of semantic and visual judgment tasks, we conducted repeated-measures ANOVAs on RT and ACC, with task type (semantic vs. visual) and difficulty (easy vs. hard) as within-subject factors. Group-level statistical results for RT and ACC are shown in Figure 2a (see Table S4 for detailed descriptive statistics). A significant main effect of difficulty was observed for RT (F_(1, 34)_ = 544.510, *p* < 0.001, η²_p_ = 0.941), with longer RTs in the hard condition compared to the easy condition, regardless of task type. Specifically, RTs were significantly longer for the semantic task in hard (1698.40 ± 50.58 ms) compared to easy condition (1296.46 ± 39.23 ms, F_(1, 34)_ = 256.09, *p* < 0.001, η^2^_p_ = 0.883), as well as for the visual task (hard: 1741.83 ± 55.46 ms; easy: 1135.54 ± 41.21 ms; F_(1, 34)_ = 536.836, *p* < 0.001, η^2^_p_ = 0.940). The main effect of task type did not reach statistical significance (F_(1, 34)_ = 3.203, *p* = 0.082). Importantly, there was a significant interaction between task type and difficulty (F_(1, 34)_ = 54.634, *p* < 0.001, η²_p_ = 0.616). Post-hoc analyses revealed that in the hard condition, there was no significant difference in RTs between the semantic and visual tasks (semantic: 1698.40 ± 50.58 ms; visual: 1741.83 ± 55.46 ms; F_(1, 34)_ = 1.138, *p* = 0.294), suggesting comparable RTs under high control demand. However, in the easy condition, RTs were significantly longer for the semantic task compared to the visual task (semantic: 1296.46 ± 39.23 ms; visual: 1135.54 ± 41.21 ms; F_(1, 34)_ = 29.456, *p* < 0.001, η²_p_ = 0.464), potentially reflecting distinct cognitive processing associated with semantic tasks. Notably, the difficulty effect (i.e., the difference between the hard and easy conditions) was smaller for semantic tasks than for visual tasks (semantic: 401.91 ± 148.51 ms; visual: 606.14 ± 154.78 ms; t_(34)_ = -7.381, *p* < 0.001). This indicates that the difficulty effect in semantic tasks captures processes more specific to semantic control, rather than reflecting general cognitive control alone. This interaction pattern supports the use of difficulty manipulations in semantic and visual tasks to define semantic control.

For ACC, a significant main effect of difficulty was observed (F_(1, 34)_ = 115.894, *p* < 0.001, η²_p_ = 0.773), with higher ACC in the easy condition compared to the hard condition (easy: 97.64 ± 0.38%; hard: 90.00 ± 0.81%). There was no difference between the difficulty effect (i.e., the difference between the hard and easy conditions) for semantic tasks and visual tasks (semantic: - 7.94 ± 5.45%; visual: -7.69 ± 5.39%; t_(34)_ = -0.215, *p* = 0.831). A significant main effect of task type was also observed (F_(1, 34)_ = 4.402, *p* = 0.043, η²_p_ = 0.115), with slightly lower ACC for the semantic task compared to the visual task (semantic: 93.13 ± 0.66%; visual: 94.51 ± 0.59%). The interaction between task type and difficulty was not significant (F_(1, 34)_ = 0.028, *p* = 0.869). These results confirm that our manipulation of control demand effectively influenced task performance. Both semantic and visual tasks showed pronounced effects of difficulty on RT and ACC, underscoring the robustness of our task design.

### 3.2 IFG, pMTG, and dmPFC showed greater activation during hard semantic task

To identify regions involved in semantic control for use as seeds in the TMS study, we localized areas showing greater activation during demanding semantic tasks compared to demanding visual tasks by contrasting the difficulty effect (Hard - Easy) in the semantic task with that in the visual task. As anticipated, this analysis revealed significant activation in the left IFG, pMTG, and dmPFC (Figure 3g, Table 1). Furthermore, ROI analysis demonstrated that the effect of semantic difficulty on activation was significantly greater than that of visual difficulty (Figure 3h–j). This robust differential activation confirmed the involvement of these regions in semantic control, justifying their selection for further investigation using TMS.

**Figure 3.**
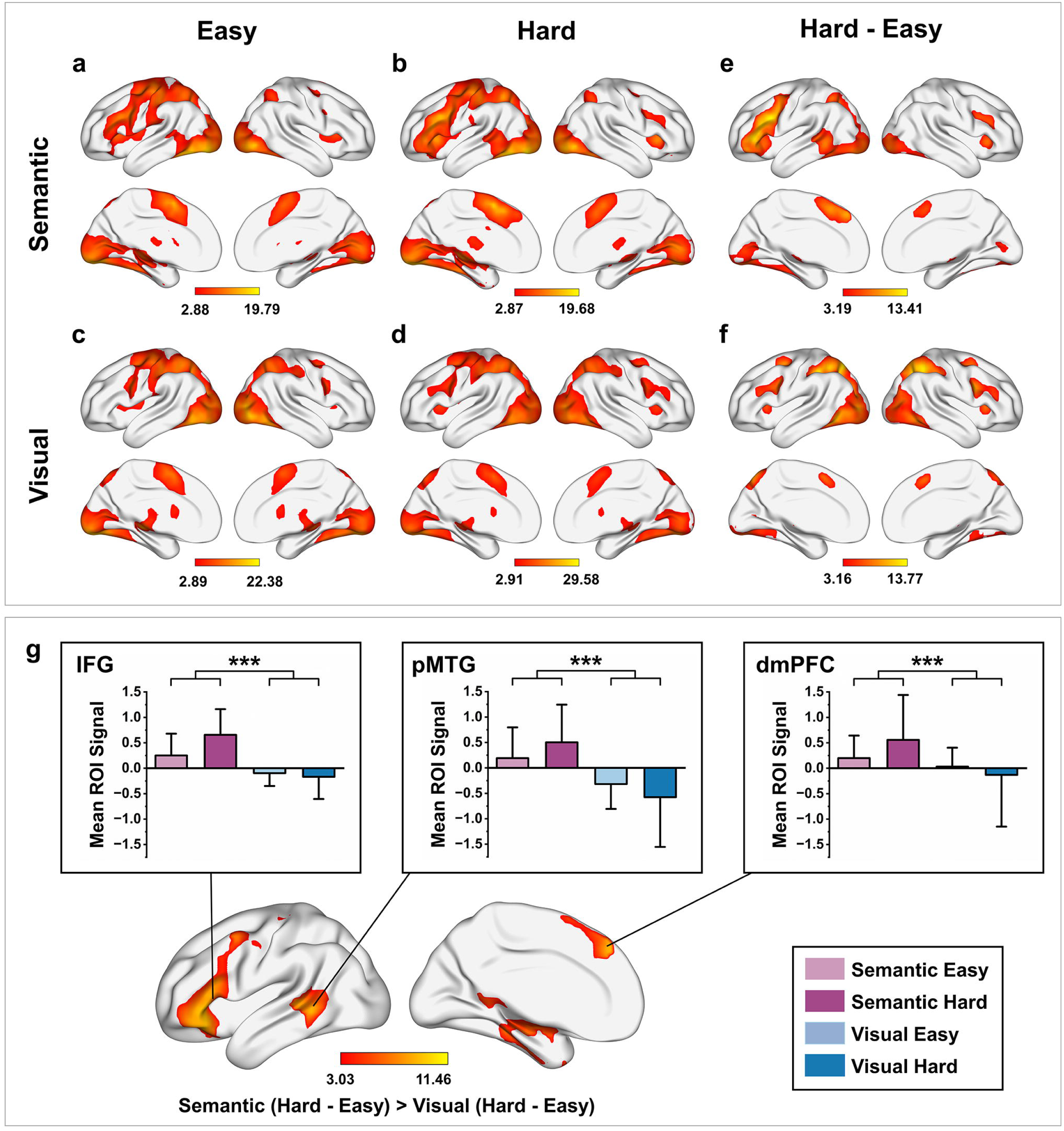
Group-level activation maps. **(a–d)** Whole-brain activation maps for each experimental condition. **(e,f)** Difficulty effects for the semantic task (Semantic Hard - Easy) and the visual task (Visual Hard - Easy). **(g)** Interaction contrast showing regions where the semantic difficulty effect exceeds the visual difficulty effect: (Semantic Hard - Easy) > (Visual Hard - Easy). A positive-activation mask derived from the contrast Semantic Hard > Rest was applied. **(h–j)** ROI analysis of the interaction contrast in the left IFG, dmPFC and pMTG. The statistical threshold for this analysis was set at *p* < 0.01, FDR-corrected at the voxel-wise level, with a minimum cluster size of 50 voxels.

**Table 1.** Brain region activations in the whole brain general linear model (GLM) analysis.

| Regions | Cluster size | T | Peak coordinates<br>(MNI) |  |  |
| --- | --- | --- | --- | --- | --- |
|  |  |  | x | y | z |
| *Semantic (Hard - Easy) > Visual (Hard - Easy)* |  |  |  |  |  |
| L IFG pars orbitalis | 2303 | 9.74 | -45 | 30 | -6 |
| L IFG pars orbitalis |  | 9.64 | -42 | 33 | -15 |
| L ITG |  | 9.38 | -42 | -12 | -27 |
| L dorsomedial prefrontal cortex | 546 | 10.39 | -9 | 39 | 42 |
| L supplementary motor areas |  | 5.81 | -6 | 12 | 66 |
| L Precuneus | 398 | 8.06 | -6 | -54 | 12 |
| L Lingual gyrus |  | 5.74 | -9 | -45 | 3 |
| L Cuneus |  | 5.48 | 0 | -87 | 21 |
| R crus II cerebellum | 348 | 11.46 | 12 | -87 | -33 |
| R crus I cerebellum |  | 8.28 | 30 | -75 | -36 |
| L pMTG | 344 | 9.44 | -63 | -45 | 0 |
| L pMTG |  | 8.75 | -54 | -36 | 3 |
| R Fusiform | 61 | 5.85 | 36 | -6 | -33 |
| R Hippocampus | 51 | 5.58 | 21 | -15 | -12 |

Beyond their general role in semantic control, all examined regions (IFG, pMTG, dmPFC) demonstrated a robust anterior–posterior functional gradient (Figure 3a–d). Anterior subregions selectively engaged during the hard semantic task, while posterior subregions responded to both easy and hard conditions. This topographical organization aligns with recent evidence that semantic control networks are anatomically proximal to the default mode network (X. Wang et al., 2024b).

### 3.3 Causal involvement of IFG, dmPFC and pMTG in Semantic Control

Previous studies have established the causal involvement of the left IFG and pMTG in semantic control; however, whether the left dmPFC also causally contributes to semantic control remains unknown. To address this critical gap, we applied TMS to these three regions. Each stimulation site was individually localized for each participant to account for anatomical and/or functional variability (see Method 2.7). To assess whether the TMS effects were specific to semantic control, we examined whether stimulation selectively impaired performance on the hard semantic task. We conducted a repeated-measures ANOVA with three within-subject factors: TMS site (IFG, pMTG, dmPFC, vertex), task type (semantic, visual), and difficulty (easy, hard). For this analysis, subdivisions within the IFG (anterior, middle) and pMTG (anterior, posterior) were treated as single regions. The analysis revealed a marginal trend toward a significant three-way interaction for RTs (F_(3,63)_ = 2.566, *p* = 0.062, η^2^_p_ = 0.109), but not for accuracy (F_(3,63)_ = 0.434, *p* = 0.729). This aligns with prior research showing that RTs are more sensitive to TMS effects than accuracy (Pobric et al., 2007; Whitney et al., 2011b). Consequently, we focused on RTs in subsequent analyses (see Table S5 for detailed descriptive statistics).

We then conducted separate two-way repeated-measures ANOVAs for semantic and visual tasks, with TMS site (IFG, pMTG, dmPFC, vertex) and difficulty (easy, hard) as within-subject factors. Critically, for semantic task RTs, we observed a significant interaction between difficulty and TMS site (F_(3,63)_ = 3.940, *p* = 0.012, η^2^_p_ = 0.158), whereas no such interaction was observed for visual task RTs (F_(3,63)_ = 0.587, *p* = 0.626). We revealed significant RT prolongations during hard semantic condition following TMS to IFG, pMTG, and, importantly, the dmPFC compared to control vertex stimulation. These effects were confirmed through post hoc comparisons using paired t-tests, with p-values corrected via Benjamini-Hochberg FDR correction (all *q* _(FDR)_ < 0.01; see Table 2 for detailed statistics). No significant RT differences were observed in easy semantic, easy visual, or hard visual conditions (all *q*_(FDR)_ > 0.05; see Table 2 for statistics), suggesting specificity to demanding semantic conditions. Furthermore, the TMS-induced RT effects (calculated as target site minus vertex differences) were significantly stronger in hard semantic condition compared to all other conditions (see Table 3 for statistics). Although the contrast between hard semantic and hard visual conditions at pMTG approached marginal significance (*q* = 0.086), the overall results strongly support the specificity of the effect and rules out the involvement of domain-general control mechanisms. Crucially, our findings replicate prior evidence of causal involvement of the left IFG and pMTG, and provide novel evidence demonstrating, for the first time, a causal role of the left dmPFC in semantic control.

**Table 2.** t- and p-values from paired T-tests comparing the TMS effect on RT between target and control site for each condition (Easy Semantic, Hard Semantic, Easy Visual, Hard Visual; N = 22)

| Target Site | Condition | $\Delta RT$ (M ± SD) | df | t | p | q(FDR) |
| --- | --- | --- | --- | --- | --- | --- |
| **IFG vs. Vertex** |  |  |  |  |  |  |
| | Easy Semantic | 73.31 ± 141.29 | 21 | 2.434 | 0.024 | 0.067 |
| | Hard Semantic | 171.391 ± 137.84 | 21 | 5.832 | 0.000 | **0.000\*\*\*** |
| | Easy Visual | 26.64 ± 98.27 | 21 | 1.272 | 0.217 | 0.289 |
| | Hard Visual | 44.74 ± 158.36 | 21 | 1.325 | 0.199 | 0.289 |
| **pMTG vs. Vertex** |  |  |  |  |  |  |
| | Easy Semantic | 50.06 ± 127.37 | 21 | 1.843 | 0.079 | 0.158 |
| | Hard Semantic | 143.49 ± 156.36 | 21 | 4.304 | 0.000 | **0.001\*\*** |
| | Easy Visual | 30.13 ± 107.40 | 21 | 1.316 | 0.202 | 0.289 |
| | Hard Visual | 55.04 ± 219.38 | 21 | 1.177 | 0.252 | 0.302 |
| **dmPFC vs. Vertex** |  |  |  |  |  |  |
| | Easy Semantic | 72.84 ± 144.29 | 21 | 2.368 | 0.028 | 0.067 |
| | Hard Semantic | 156.29 ± 164.19 | 21 | 4.465 | 0.000 | **0.001\*\*** |
| | Easy Visual | 19.11 ± 96.89 | 21 | 0.925 | 0.365 | 0.398 |
| | Hard Visual | 8.56 ± 211.33 | 21 | 0.190 | 0.851 | 0.851 |

Note. The table presents both uncorrected p-values and Benjamini-Hochberg FDR-adjusted q- values. Significance thresholds were defined as: q < 0.01 (\*\*) and q < 0.001 (\*\*\*).

**Table 3.** t- and p-values from paired T-tests comparing the TMS effect on RT between Hard Semantic condition and other conditions at each target sites (IFG, pMTG, dmPFC; N = 22)

| | Target Site | $\Delta RT$ (M ± SD) | df | t | p | q<sub>(FDR)</sub> |
| --- | --- | --- | --- | --- | --- | --- |
| **Semantic Hard vs. Semantic Easy** |  |  |  |  |  |  |
| | IFG | 98.08 ± 117.53 | 21 | 2.591 | 0.017 | **0.021\*** |
| | pMTG | 93.41 ± 148.45 | 21 | 2.951 | 0.008 | **0.011\*** |
| | dmPFC | 83.45 ± 153.29 | 21 | 2.554 | 0.018 | **0.021\*** |
| **Semantic Hard vs. Visual Easy** |  |  |  |  |  |  |
| | IFG | 114.75 ± 148.33 | 21 | 4.577 | 0.000 | **0.001\*\*** |
| | pMTG | 113.33 ± 173.28 | 21 | 3.068 | 0.006 | **0.010\*** |
| | dmPFC | 137.18 ± 168.11 | 21 | 3.827 | 0.001 | **0.004\*\*** |
| **Semantic Hard vs. Visual Hard** |  |  |  |  |  |  |
| | IFG | 126.65 ± 185.73 | 21 | 3.198 | 0.004 | **0.010\*\*** |
| | pMTG | 88.43 ± 230.03 | 21 | 1.803 | 0.086 | 0.086 |
| | dmPFC | 147.74 ± 203.89 | 21 | 3.399 | 0.003 | **0.008\*\*** |

Note. The table presents both uncorrected p-values and Benjamini-Hochberg FDR-adjusted q-
values. Significance thresholds were defined as: q < 0.05 (\*) and q < 0.01 (\*\*).

### 3.4 Causal evidence of functional differentiation within IFG and pMTG

Since we observed a robust anterior–posterior functional gradient in IFG, pMTG and dmPFC in task fMRI, with anterior subregions selectively being engaged during the hard semantic task, while posterior subregions responding to both easy and hard conditions. To further explore functional differentiation within the left IFG and pMTG, participants were divided based on stimulation sites (see Method 2.8). Given that all dmPFC stimulation sites were located within the anterior portion, this region was not further subdivided. We examined whether TMS of specific subregions affected behavior differently compared to Vertex stimulation across task conditions by conducting paired t-tests with permutation tests and Benjamini-Hochberg FDR correction. The locations of the TMS stimulation sites for each subregion group within the left IFG and pMTG, as well as the behavioral effects on RT for each task condition, are shown in shown in Figure 4. Subregion-specific behavioral effects were observed within both the left IFG and pMTG (see Table S7 for detailed statistics). Stimulation of the a-IFG resulted in significantly longer RTs under hard semantic condition compared to Vertex stimulation (t_(6)_ = 4.615, *q*_(FDR)_ = 0.049), whereas no significant differences were found under the other conditions (all *q*_(FDR)_ > 0.05). In contrast, stimulation of the mid-IFG led to significantly longer RTs under both the easy (t_(15)_ = 2.764,*q*_(FDR)_ = 0.049) and hard semantic conditions (t_(15)_ = 4.386,*q*_(FDR)_ = 0.010) compared to Vertex stimulation. For the pMTG, stimulation of the a-pMTG resulted in significantly longer RTs under both the hard semantic (t_(9)_ = 4.723,*q*_(FDR)_ = 0.016) and hard visual (t_(11)_ = 3.281,*q*_(FDR)_ = 0.031) conditions, compared to Vertex stimulation, while no significant differences were observed under the easy conditions (all *q*_(FDR)_ > 0.05). Stimulation of the p-pMTG did not produce significant effects under any condition (all *q*_(FDR)_ > 0.05). Notably, no significant behavioral effects were observed in the ACC following stimulation of different subregions within the left IFG and pMTG under any task condition (see Figure S1 and Table S8 for detailed statistics). These findings highlight the functional differentiation within each region, with the anterior subregions of the left IFG and pMTG playing a more critical role in demanding semantic tasks.

**Figure 4.**
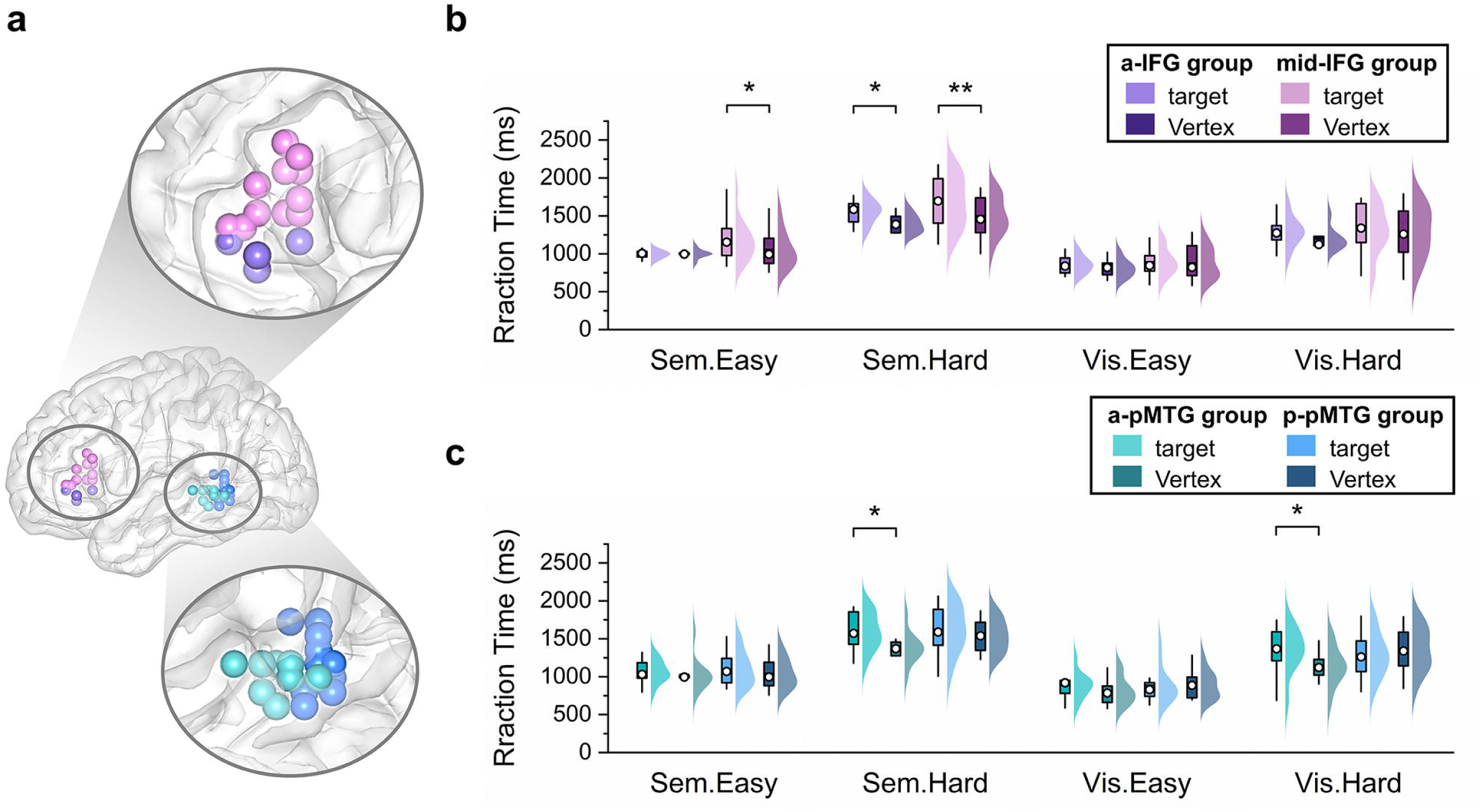
RT differences in the TMS experiment between stimulation on the anterior and posterior subregions of the IFG and pMTG compared to Vertex stimulation. **(a)** Locations of TMS stimulation site are visualized on the standard ICBM152 template surface, with each site represented by a color-coded dot for each subject: aIFG in purple, mid-IFG in pink, a-pMTG in teal, and p-pMTG in blue. **(b)** RT results for subregions of IFG under each task conditions, with participants divided into two groups based on stimulation site (a- and mid-IFG), and Vertex stimulation of the same group participants used as a comparison. **(c)** RT results for subregions of pMTG under each task conditions, with participants divided into two groups based on stimulation site (a- and p-pMTG), with Vertex stimulation of the same group used as a comparison. The x-axis displays the four conditions, while the y-axis shows mean RT of each condition. Boxplots show the median (denoted by hollow circles) and 1.5 times the interquartile range. Half-violin plots demonstrate the data distribution, with each region represented by a color matching the TMS stimulation site and corresponding Vertex represented by a darker color. Abbreviations: a-IFG, the anterior subregion of inferior frontal gyrus; mid-IFG, the middle subregion of inferior frontal gyrus; a-pMTG, the anterior subregion of posterior temporal gyrus; p-pMTG, the posterior subregion of posterior temporal gyrus; RT, reaction time. Statistical significance thresholds were defined as follows: *q*_(FDR)_ < 0.05 (*) and *q*_(FDR)_ < 0.01(**).

### 3.5 Coordinated contribution of IFG, pMTG, and dmPFC to semantic control

After establishing the causal involvement of the left IFG, pMTG, and dmPFC in semantic control, we further investigated whether these regions function collectively or independently. Specifically, we tested whether the combined activation patterns of these regions better predicted the behavioral differences between hard and easy semantic tasks compared to the activation patterns of any single region. Our analysis showed that the combined activation patterns of the left IFG, pMTG, and dmPFC significantly predicted the behavioral differences in RT between hard and easy semantic tasks (*r* = 0.51, *p* = 0.001, permutation tests with 1000 iterations; Figure 5a). In contrast, prediction models based on the activation patterns of individual regions were not significant (IFG: *r* = 0.29, *p* = 0.203; pMTG: *r* = 0.24, *p* = 0.332; dmPFC: *r* = 0.25, *p* = 0.281; all p-values derived from permutation tests with 1000 iterations; Figure 5b). Direct comparisons of mean prediction errors (MAE and MSE) between the combined model and single-region models showed no significant differences (all ps > 0.05), indicating comparable point-prediction accuracy among the four models. However, prediction errors alone do not account for differences in model complexity and parameter uncertainty. To explicitly balance these factors, we conducted Bayesian model selection. Higher model evidence indicates a superior balance between model complexity and fit quality, thus providing stronger explanatory power. According to the criteria established by Kass & Raftery (1995), a logBF greater than 5 constitutes decisive evidence. Accordingly, our results provided decisive evidence favoring the combined model over all single-region models (combined vs. IFG: logBF = 12.79; combined vs. dmPFC: logBF = 9.87; combined vs. pMTG: logBF = 11.10). These converging results strongly suggest that the left IFG, pMTG, and dmPFC operate in a coordinated manner to support semantic control, rather than functioning independently.

**Figure 5.**
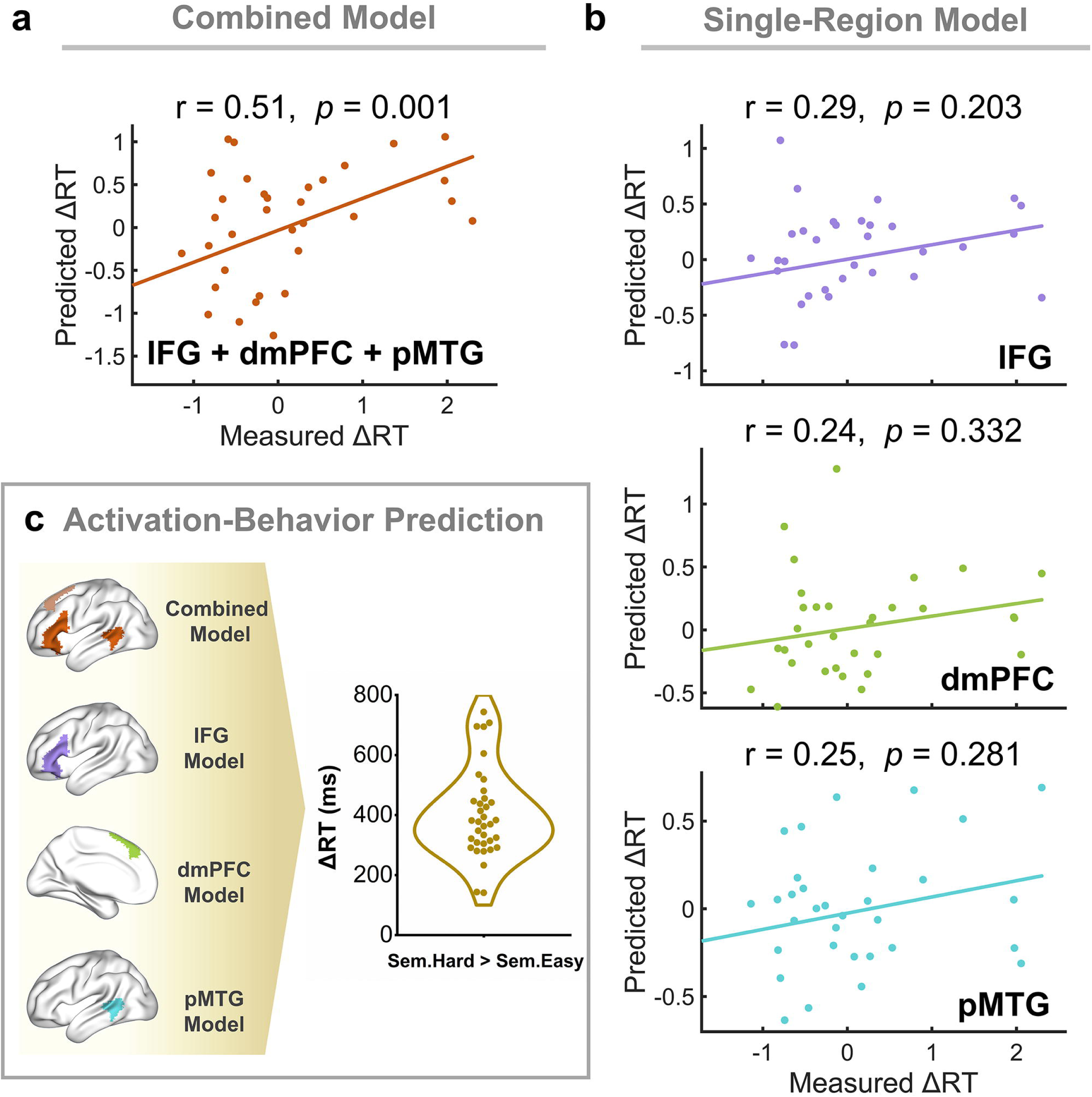
Accuracy and significance of the multivariate machine learning-based (RVR-based) prediction models. **(a)** The combined model predicting semantic difficulty-induced reaction time changes (ΔRT_Sem.Hard-Sem.Easy_) using integrated activation patterns from the left IFG, dmPFC, and pMTG. This model leverages the synergistic contributions of these regions to capture behavioral variations associated with task difficulty. **(b)** Single-region models predicting ΔRT_Sem.Hard-Sem.Easy_ based on activation patterns derived from IFG-, dmPFC-, and pMTG-specific analyses individually. **(c)** Schematic illustrates the activation-behavior prediction methodology. The brain maps in blue denote regions used for feature extraction in combined and single-region models. Only voxels within these blue region masks were selected for extraction of beta values as predictive features for each participant, with ΔRT_Sem.Hard-Sem.Easy_ as the target behavioral measure. The R value represents the Pearson correlation coefficient between predicted and actual values, and model significance and P-values were determined using 1,000 permutation tests.

### 3.6 Modulation of connectivity by semantic control

To further investigate how the IFG, dmPFC, and pMTG interact to support semantic control, we employed DCM to examine the effective connectivity between these regions. This analysis allowed us to explore the directional influences between regions and how semantic task difficulty modulates their connectivity. Semantic control significantly modulated the autoinhibition of individual regions and the connectivity between them: Hard semantic condition reduced autoinhibition in IFG and pMTG, as indicated by negative values (-2.03 and -1.14), making them more responsive to input from the network. In contrast, the hard semantic condition increased autoinhibition in dmPFC, as indicated by a positive value (1.91), rendering it less responsive to network inputs (Figure 6b). Semantic control also altered the nature of connectivity between regions, transforming excitatory influences (positive values) into inhibitory influences (negative values) and vice versa (Figure 6b). Specifically, during the hard semantic condition, IFG and pMTG exhibited reciprocal excitatory influences (Figure 6b), whereas they showed reciprocal inhibitory influences during the easy semantic task (Figure 6a). Similarly, IFG and dmPFC demonstrated reciprocal inhibitory influences during the hard condition, contrasting with their reciprocal excitatory influences during the easy semantic task. For the connection between pMTG and dmPFC, semantic control resulted in an inhibitory influence during the hard condition but an excitatory influence during the easy semantic task. Conversely, dmPFC and pMTG showed excitatory influences during the hard condition, whereas their interaction was inhibitory during the easier semantic task (Figure 6a).

**Figure 6.**
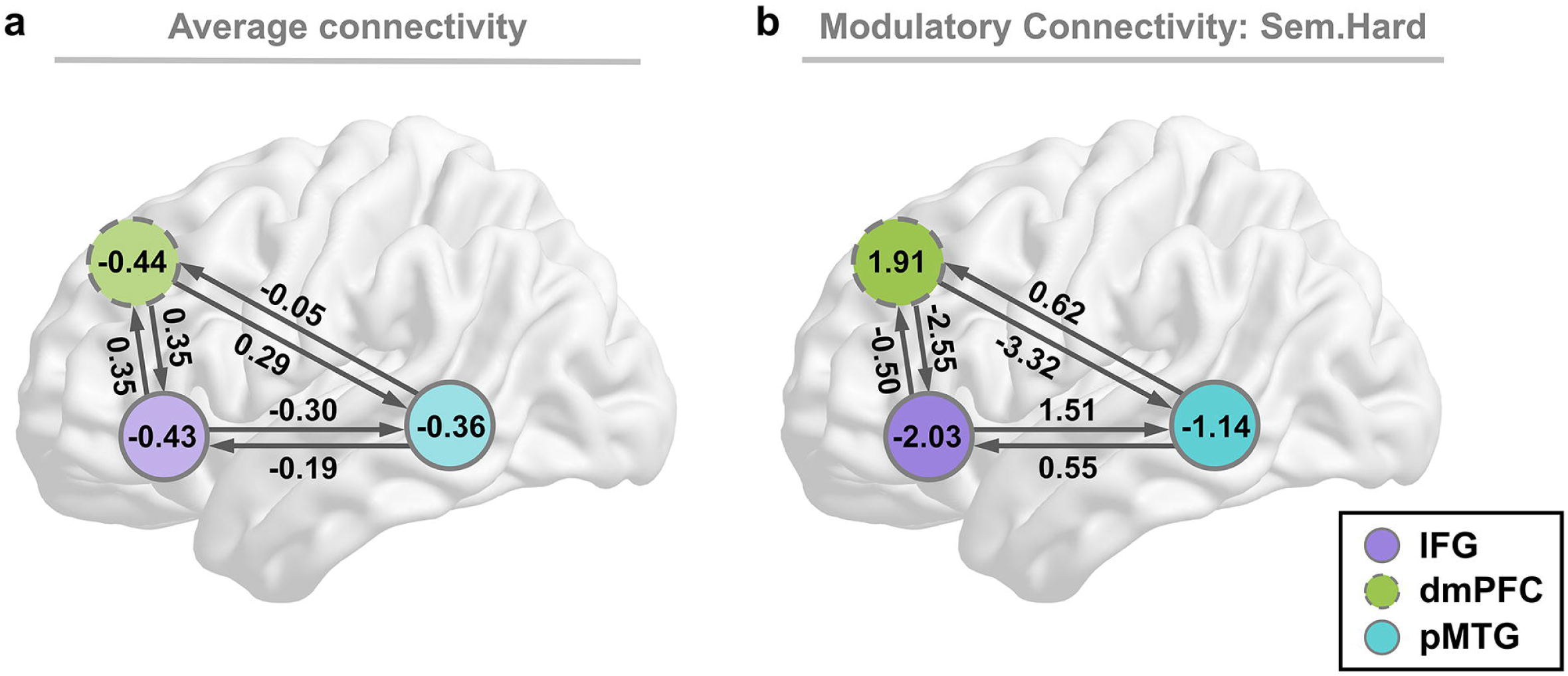
DCM results. **(a)** Matrix A representing the average effective connectivity across all semantic conditions (Easy and Hard). The parameters within circle denote autoinhibition connectivity, while parameters shown on the lines indicate connectivity between regions. **(b)** Matrix B illustrating the modulatory effect on effective connectivity between regions and autoinhibition under the hard semantic condition. Positive effective connectivity values indicate excitatory connectivity, while negative values reflect inhibitory connectivity. Parameters with stronger evidence (posterior probability > 95%) are presented and subthreshold parameters are marked with “n.s.”.

### 3.7 Associations between semantic control performance and connectivity parameters

To investigate how effective connectivity varies in response to behavioral variability in semantic control demand, we correlated effective connectivity parameters from DCM with individual behavioral variability in the residual IE score (see section 2.6.4). We found that reduced autoinhibition (i.e., more negative connectivity values) in the left IFG was associated with poorer performance (higher RV score) across all semantic conditions (Figure S2a). As the semantic tasks became more challenging, increased autoinhibition in the left dmPFC was linked to poorer performance, whereas reduced autoinhibition in the left pMTG correlated with poorer performance (Figure S2b). These findings suggest that as semantic tasks become more difficult, there is a need of greater involvement of the semantic control network to support task performance.

## 4 Discussion

The current study disentangles the causal roles and dynamic interactions among the left IFG, dmPFC, and pMTG in semantic control. By combining task-fMRI localization with individualized TMS, we replicated prior evidence supporting the causal involvement of the left IFG and pMTG, and, critically, provided the first direct causal evidence implicating the dmPFC in semantic control. Stimulation-site comparisons also revealed anterior–posterior functional heterogeneity in the IFG and pMTG, supporting systematic topographical organization. Multivariate prediction analyses further showed that combined activity across these nodes predicted performance better than any single-region model. DCM analyses revealed that semantic difficulty bidirectionally modulates interregional interactions, indicating dynamic network reconfiguration. Together, these findings advance our mechanistic understanding of semantic control as a topographically organized, dynamically reconfiguring network.

### 4.1 The left IFG, pMTG, and dmPFC—key regions for semantic control

Our results provide new causal evidence for the crucial roles of the left IFG, dmPFC, and pMTG in semantic control. While substantial NIBS studies have established the necessity of the left IFG and pMTG in this domain (Whitney et al., 2011b, 2012; Davey et al., 2015; Hallam et al., 2016; Teige et al., 2018; Alonso et al., 2024), prior research has rarely emphasized the left dmPFC’s contribution. Anatomically, the left dmPFC serves as a “connector” hub at the intersection of multiple large-scale networks, including MDN, SCN, and DMN (Roger et al., 2022; Chiou et al., 2023; X. Wang et al., 2024a), which has sometimes led to misinterpretations of its role in semantic tasks. Recent evidence, including our findings, highlights the dmPFC’s critical involvement in semantic control (Binder & Desai, 2011; Jackson, 2021; Graessner et al., 2021; Heo et al., 2023; X. Wang et al., 2024a). TMS-induced disruption of these three regions selectively impaired performance on hard semantic tasks without affecting non-semantic tasks of comparable difficulty, providing the first direct evidence for the necessity of the left dmPFC in semantic control, while replicating prior findings of causal involvement for the left IFG and pMTG. Collectively, these results highlight a distributed network comprising the left IFG, pMTG, and dmPFC as critical for semantic control.

### 4.2 Functional heterogeneity within key regions in semantic control

Another critical insight from this study is the functional heterogeneity observed within the left IFG and pMTG. TMS disruption targeting anterior versus middle/posterior subregions in these regions revealed distinct behavioral profiles. In the left IFG, inhibition of the anterior subregion (approximating the pars orbitalis) selectively impaired performance under high semantic control demands, whereas inhibition of the middle subregion (pars triangularis) impaired performance across both easy and hard semantic tasks. This pattern supports prior fMRI studies demonstrating functional differentiation within the IFG (J. Wang et al., 2020; Fedorenko & Blank, 2020; M. Zhang et al., 2022; Diveica et al., 2023) and is consistent with Badre’s two-process model, which posits that the anterior IFG is more specialized for controlled access to stored conceptual representations, while the posterior IFG is more involved in selection process that operates post-retrieval (Badre, 2005; Badre & Wagner, 2007).

A similar pattern of functional heterogeneity was observed in the left pMTG. TMS inhibition of the anterior pMTG selectively impaired performance under high control demand conditions, whereas inhibiting the posterior pMTG produced no significant behavioral effects. Converging structural and resting-state functional connectivity evidence indicates that the left MTG can be parcellated into four subregions (Xu et al., 2015, 2019), two of which correspond to the anterior and posterior pMTG sites targeted in our study. The anterior subregion preferentially connects with left IFG (pars orbital), medial middle frontal gyrus (MFG), angular gyrus (AG) and medial superior frontal gyrus (SFG)—regions implicated in controlled semantic retrieval and higher-order cognition—whereas the posterior subregion shows stronger coupling with bilateral MTG involved in action and object-observation processes (Xu et al., 2019). This connectivity profile parsimoniously explains our behavioral dissociation. Consistent with this distinction, recent work proposes that the pMTG functions as a multimodal integration hub, synthesizing information from diverse sensory or action-processing pathways to enrich perceptual detail and associative information for semantic representation or retrieval (Kuhnke et al., 2023). Cross-domain meta-analytic findings further reveal that posterior pMTG is activated both during semantic control and tool-related tasks (Hodgson et al., 2022), supporting a broader multimodal representational role. Together, these findings position the anterior pMTG as a critical node within the semantic control circuit, whose transient disruption reliably impaired performance under high control demands, whereas perturbing the posterior pMTG yielded no measurable deficit.

These findings may reflect adaptations to the computational demands of cross-network integration. Recent theoretical frameworks propose a hybrid neurocognitive architecture, in which complex cognitive abilities emerge from dynamic interactions across multiple functional networks (Herbet & Duffau, 2020). Within this framework, certain regions—termed “connector hubs”—integrate and coordinate information from diverse networks in response to changing task demands, thereby facilitating cognitive flexibility (Roger et al., 2022). Semantic task performance relies on interactions among the SCN, DMN, and MDN (Davey et al., 2016; Evans et al., 2020; X. Wang et al., 2021). The DMN—including regions such as AG and ATL—supports the automatic activation of stored semantic representations and the maintenance of semantic goals (Patterson et al., 2007; Binder et al., 2009; Seghier et al., 2010; Humphreys et al., 2015; X. Wang et al., 2021), whereas the MDN implements executive control processes aligned with current task goals (Duncan, 2010; Badre & Nee, 2018; Assem et al., 2020). SCN regions occupy an intermediate spatial and functional position between these two systems (Chiou et al., 2023; X. Wang et al., 2024a), serving as “connector” hubs that mediate the selective transfer of information from semantic memory systems to executive control networks during demanding semantic tasks (Davey et al., 2016; X. Wang et al., 2020; Roger et al., 2022). Functional subdivisions within SCN regions may facilitate this process by mitigating potential interference from globally interconnected networks and promote efficient information exchange (X. Wang et al., 2024b). One possibility is that such heterogeneity reflects adaptations to the computational demands of cross-network integration, optimizing flexible semantic control.

### 4.3 Functional interactions of key regions in the SCN

Our findings provide converging evidence that the left IFG, pMTG, and dmPFC interact coordinately during controlled semantic retrieval. First, a combined predictive model using activation patterns from all three regions outperformed single-region models, highlighting their synergistic contributions and supporting the view that flexible semantic retrieval relies on a distributed control network (Lambon Ralph et al., 2017; Hodgson et al., 2021). Second, DCM analyses further revealed that effective connectivity among these regions varies with semantic control demands. Under hard semantic conditions, the left IFG and pMTG exhibited strong reciprocal excitation, with stronger top-down modulation from the IFG. This may suggest that the IFG may regulate retrieval processes, while the pMTG feeds retrieved information back to refine or update strategies. Meanwhile, the left dmPFC exerted inhibitory effects on both the IFG and pMTG, likely serving as a regulatory “brake” to prevent overactivation. The inhibitory influence of IFG on dmPFC may serve as a feedback pathway to help maintain network balance by updating contextual signals and adjusting the intensity of top-down control in response to task demands (Badre & Nee, 2018). Although these interpretations are based on effective connectivity rather than direct neurophysiological evidence, they strongly support the view that SCN regions dynamically modulate control intensity through reciprocal interactions, enabling the network to flexibly handle escalating semantic control demands while maintaining both flexibility and stability (Cole et al., 2013; J. Y. Jung et al., 2021; X. Wang et al., 2024a).

Our results extend previous findings while revealing key differences. Hallam et al. (2016) combined TMS and fMRI and found that inhibiting the left IFG increased activation in the pMTG and pre-SMA during difficult semantic tasks, despite no significant behavioral impairments. This compensatory activation pattern suggests that suppression of one SCN node may trigger upregulation in others, supporting a cooperative network view. By contrast, Hartwigsen et al. (2017) examined effective connectivity during a sentence integration task and found that the left IFG exerted stronger inhibitory influence on the pMTG when sentences contained anomalous endings. This implies that under semantic conflict, the left IFG suppresses activations in temporal regions to prevent interference with sentence-level integration (Hartwigsen et al., 2017). In contrast, our hard semantic task emphasized strengthening weak associations between concepts, requiring the enhancement rather than suppression of semantic activations. This distinction highlights that SCN connectivity patterns are not only task-dependent but also critically shaped by the nature of the underlying processing operations—whether semantic activation must be suppressed or amplified to support successful task performance. Moreover, previous studies largely overlooked the role of the left dmPFC. By incorporating this region, our findings revealed semantic control demand-sensitive interactions between the dmPFC, IFG, and pMTG, offering a more comprehensive view of the SCN’s adaptive functional organization during controlled semantic retrieval.

### 4.4 Limitation

Several limitations should be noted. First, while TMS inhibition of the left IFG, pMTG, and dmPFC impaired performance, its broader effects on the semantic control network remain unclear. Given that TMS can influence connected regions beyond the target (Bergmann et al., 2021), it is uncertain whether observed deficits reflect local disruption or network-wide disturbances. Combining TMS with functional imaging could address this limitation. Second, while our results suggest functional heterogeneity within the left IFG and pMTG, the small sample size limits statistical power. The functional organization of the IFG in semantic control remains debated (Thompson-Schill et al., 1997; Badre, 2005; Thompson-Schill & Botvinick, 2006; Martin & Cheng, 2006; Snyder et al., 2011), and larger studies with finer targeting are needed to validate these findings. Third, we focused on SCN regions under high semantic control demands, but did not examine how the SCN collaborates with broader networks such as the DMN and MDN, which are critical for flexible semantic processing (Jackson et al., 2021; J. Jung & Lambon Ralph, 2023; Branzi & Lambon Ralph, 2023; X. Wang et al., 2024a). Future investigations should examine how these systems collaboratively support complex semantic tasks, providing a more comprehensive understanding of the neural mechanisms underlying semantic control. Finally, the limited temporal resolution of fMRI highlights the need for techniques such as intracranial EEG to map the fine-grained temporal dynamics of semantic control.

## 5. Conclusion

In conclusion, this study demonstrates the causal involvement, functional heterogeneity, and effective interaction patterns of the key regions in SCN. Specifically, the anterior portions of the left IFG and pMTG, rather than their posterior parts, are more causally involved in semantic control. Moreover, these regions coordinated through finely tuned excitatory and inhibitory influences under high semantic control demands, supporting flexible semantic retrieval and manipulation of semantic information. Collectively, these findings significantly advance our understanding of the neural mechanisms underlying semantic control and provide a more nuanced framework for exploring how the brain achieves semantic flexibility and efficiency in response to varying task demands.

## Acknowledgments

The study is supported by the National Social Science Foundation of China (No. 20&ZD296), the Key-Area Research and Development Program of Guangdong Province (No. 2019B030335001), and the National Natural Science Foundation of China (No.32100889 and No.32300881), Research Center for Brain Cognition and Human Development, Guangdong, China (No. 2024B0303390003). We are grateful to Xinling Wang for providing guidance on the design and piloting of the TMS experiment, and to Jiexian Gong for assistance in developing the stimulus presentation scripts.

