## Supplemental Data 1 for "Mapping the functional differentiation and interactions among the inferior, medial frontal and posterior temporal cortex in semantic control"

**Table S1. Psycholinguistic variables for each condition across six sessions (Mean and SD values)**

| Session | Condition | Association strength | stroke | frequency (logW) | familiarity | imageability |
| --- | --- | --- | --- | --- | --- | --- |
| fMRI task | Sem.Easy | 0.82 ± 0.5 | 16.66 (4.91) | 2.48 (0.57) | 6.08 (0.16) | 6.13 (0.15) |
|  | Sem.Hard | 0.45 (0.10) | 16.43 (4.69) | 2.41 (0.58) | 6.07 (0.17) | 6.10 (0.16) |
| TMS task 1 | Sem.Easy | 0.81 (0.08) | 17.15 (4.60) | 2.37 (0.52) | 6.08 (0.17) | 6.10 (0.18) |
|  | Sem.Hard | 0.44 (0.11) | 16.82 (4.77) | 2.38 (0.53) | 6.11 (0.17) | 6.10 (0.17) |
| TMS task 2 | Sem.Easy | 0.83 (0.06) | 17.03 (4.93) | 2.39 (0.49) | 6.08 (0.17) | 6.11 (0.17) |
|  | Sem.Hard | 0.47 (0.09) | 17.25 (4.58) | 2.39 (0.51) | 6.11 (0.18) | 6.10 (0.16) |
| TMS task 3 | Sem.Easy | 0.82 (0.06) | 17.00 (5.03) | 2.73 (0.50) | 6.11 (0.17) | 6.10 (0.17) |
|  | Sem.Hard | 0.47 (0.10) | 17.03 (4.72) | 2.39 (0.50) | 6.10 (0.17) | 6.11 (0.18) |
| TMS task 4 | Sem.Easy | 0.82 (0.06) | 17.22 (4.88) | 2.40 (0.51) | 6.09 (0.19) | 6.12 (0.18) |
|  | Sem.Hard | 0.44 (0.10) | 16.96 (5.08) | 2.38 (0.49) | 6.10 (0.17) | 6.09 (0.17) |
| TMS task 5 | Sem.Easy | 0.83 (0.06) | 17.06 (5.04) | 2.39 (0.51) | 6.10 (0.16) | 6.11 (0.17) |
|  | Sem.Hard | 0.45 (0.13) | 17.13 (4.69) | 2.40 (0.51) | 6.09 (0.17) | 6.12 (0.18) |

**Table S2. Independent t-test results for association strength between conditions in fMRI and TMS Tasks**

| Session | Condition | Mean | SD | t | p |
| --- | --- | --- | --- | --- | --- |
| fMRI task | Sem.Easy | 0.82 | 0.05 | 24.40 | 0.000 |
|  | Sem.Hard | 0.45 | 0.10 |  |  |
| TMS task 1 | Sem.Easy | 0.81 | 0.08 | 21.47 | 0.000 |
|  | Sem.Hard | 0.44 | 0.11 |  |  |
| TMS task 2 | Sem.Easy | 0.83 | 0.06 | 25.81 | 0.000 |
|  | Sem.Hard | 0.47 | 0.09 |  |  |
| TMS task 3 | Sem.Easy | 0.82 | 0.06 | 24.39 | 0.002 |
|  | Sem.Hard | 0.47 | 0.10 |  |  |
| TMS task 4 | Sem.Easy | 0.82 | 0.06 | 24.84 | 0.000 |
|  | Sem.Hard | 0.44 | 0.10 |  |  |
| TMS task 5 | Sem.Easy | 0.83 | 0.06 | 21.50 | 0.000 |
|  | Sem.Hard | 0.45 | 0.13 |  |  |

**Table S3. Two-way ANOVA results for psycholinguistic variables across conditions and sessions**

|  | Source | df | F | Sig. |
| --- | --- | --- | --- | --- |
| Stroke | Condition | 1 | 0.20 | 0.657 |
|  | Session | 5 | 0.92 | 0.469 |
|  | Condition ×Session | 5 | 0.24 | 0.944 |
| Frequency (logW) | Condition | 1 | 0.14 | 0.709 |
|  | Session | 5 | 0.97 | 0.437 |
|  | Condition ×Session | 5 | 0.49 | 0.786 |
| Familiarity | Condition | 1 | 1.2 | 0.27 |
|  | Session | 5 | 1.6 | 0.15 |
|  | Condition ×Session | 5 | 1.8 | 0.10 |
| Imageability | Condition | 1 | 1.29 | 0.257 |
|  | Session | 5 | 0.72 | 0.610 |
|  | Condition ×Session | 5 | 1.48 | 0.230 |

**Table S4. The average RT (ms) and ACC (%) from fMRI task across four conditions (Mean and SD values)**

| Measure | Sem.Easy | Sem.Hard | Vis.Easy | Vis.Hard |
| --- | --- | --- | --- | --- |
| RT | 1296.46 (232.06) | 1698.40 (299.25) | 1135.54 (243.81) | 1741.83 (328.12) |
| ACC | 97.00 (2.84) | 89.26 (6.20) | 98.29 (2.12) | 90.74 (5.72) |

**Table S5. The average RT (ms) and ACC (%) for four TMS stimulation regions across four conditions (Mean and SD values)**

| Measure | Condition | IFG | dmPFC | pMTG | Vertex |
| --- | --- | --- | --- | --- | --- |
| RT | Sem.Easy | 1110.49 (228.59) | 1110.02 (202.277) | 1087.24 (193.13) | 1037.18 (201.43) |
|  | Sem.Hard | 1638.06 (292.41) | 1622.96 (277.14) | 1610.14 (281.74) | 1466.67 (225.35) |
|  | Vis.Easy | 883.59 (160.07) | 876.06 (172.73) | 887.09 (209.18) | 856.95 (184.81) |
|  | Vis.Hard | 1305.03 (278.01) | 1268.85 (261.34) | 1315.33 (286.95) | 1260.29 (292.24) |
| ACC | Sem.Easy | 98.03 (2.03) | 97.42 (2.51) | 96.74 (3.39) | 96.29 (3.04) |
|  | Sem.Hard | 89.24 (6.99) | 88.79 (5.37) | 87.58 (6.97) | 87.35 (7.86) |
|  | Vis.Easy | 97.58 (2.71) | 97.95 (2.12) | 98.41 (2.21) | 96.82 (3.41) |
|  | Vis.Hard | 93.64 (4.07) | 92.27 (4.20) | 92.80 (4.94) | 92.27 (4.81) |

**Table S6. The average RT (ms) and ACC (%) for TMS stimulation subregions of IFG and pMTG across four conditions (Mean and SD values)**

| Measure | Subregion | Site | Sem.Easy | Sem.Hard | Vis.Easy | Vis.Hard |
| --- | --- | --- | --- | --- | --- | --- |
| RT | a-IFG group | target | 991.90 (51.23) | 1550.94 (143.62) | 858.35 (110.92) | 1294.93 (187.91) |
|  |  | Vertex | 998.13 (57.62) | 1395.27(113.49) | 808.85 (111.94) | 1201.72 (139.22) |
|  | mid-IFG group | target | 1165.83 (249.61) | 1678.71 (323.85) | 895.38(172.32) | 1309.75 (302.75) |
|  |  | Vertex | 1055.40 (232.83) | 1499.99 (248.18) | 879.40 (200.96) | 1287.62 (328.89) |
|  | a-pMG group | target | 1070.58 (143.53) | 1595.23 (231.44) | 885.53(167.07) | 1347.34 (281.45) |
|  |  | Vertex | 1030.63(203.79) | 1378.91 (198.49) | 814.06 (172.09) | 1139.36 (242.39) |
|  | p-pMG group | target | 1101.11 (218.37) | 1622.56 (306.48) | 888.38(230.89) | 1288.66(276.62) |
|  |  | Vertex | 1042.63(190.60) | 1539.80 (210.41) | 892.69 (179.67) | 1361.06 (279.56) |
| ACC | a-IFG group | target | 98.81 (1.47) | 92.14(3.85) | 97.38 (2.80) | 92.38(2.16) |
|  |  | Vertex | 97.38 (2.50) | 83.33(7.87) | 97.86 (2.13) | 92.62 (4.16) |
|  | mid-IFG group | target | 97.67 (2.09) | 87.89 (7.47) | 97.67(2.57) | 94.22 (4.47) |
|  |  | Vertex | 95.78 (3.03) | 89.22 (6.83) | 96.33(3.66) | 92.11 (4.92) |
|  | a-pMG group | target | 96.50 (2.29) | 87.00 (5.67) | 98.17 (1.57) | 93.67 (3.14) |
|  |  | Vertex | 96.17 (2.79) | 84.00 (8.60) | 98.00 (1.80) | 90.67 (4.42) |
|  | p-pMG group | target | 96.94 (3.96) | 88.06 (7.60) | 98.61 (2.53) | 92.08 (5.78) |
|  |  | Vertex | 96.39(3.11) | 90.14 (5.42) | 95.83 (3.94) | 93.61 (4.50) |

**Table S7 t- and p-values from paired T-tests comparing** **the TMS effect on RT (ms) between target site and correspondent control Vertex site at each subregion group (a-IFG, mid-IFG, a-pMTG, p-pMTG)**

|  | **Condition** | **ΔRT (M ± SD)** | **df** | **t** | ***p*_(Perm)_** | ***q*_(FDR)_** |
| --- | --- | --- | --- | --- | --- | --- |
| **a-IFG vs. Vertex** |  |  |  |  |  |  |
| (N = 7) | Sem.Easy | -6.24 ± 56.11 | 6 | -0.29 | 0.769 | 0.821 |
|  | Sem.Hard | 155.67 ± 89.24 | 6 | 4.62 | 0.015 | **0.049*** |
|  | Vis.Easy | 49.50 ± 58.33 | 6 | 2.25 | 0.083 | 0.166 |
|  | Vis.Hard | 93.21 ± 132.27 | 6 | 1.86 | 0.152 | 0.221 |
| **mid-IFG vs. Vertex** |  |  |  |  |  |  |
| (N = 15) | Sem.Easy | 110.44 ± 154.77 | 14 | 2.76 | 0.015 | **0.049*** |
|  | Sem.Hard | 178.73 ± 157.81 | 14 | 4.39 | 0.001 | **0.010**** |
|  | Vis.Easy | 15.98 ± 112.45 | 14 | 0.55 | 0.585 | 0.704 |
|  | Vis.Hard | 21.12 ± 168.51 | 14 | 0.51 | 0.616 | 0.704 |
| **a-pMTG vs. Vertex** |  |  |  |  |  |  |
| (N = 10) | Sem.Easy | 39.96 ± 141.91 | 9 | 0.89 | 0.379 | 0.506 |
|  | Sem.Hard | 216.32 ± 114.85 | 9 | 4.72 | 0.002 | **0.016*** |
|  | Vis.Easy | 71.47 ± 109.72 | 9 | 2.06 | 0.075 | 0.166 |
|  | Vis.Hard | 207.97 ± 200.45 | 9 | 3.28 | 0.006 | **0.031*** |
| **p-pMTG vs. Vertex** |  |  |  |  |  |  |
| (N = 12) | Sem.Easy | 58.48 ± 119.69 | 11 | 1.69 | 0.114 | 0.186 |
|  | Sem.Hard | 82.76 ± 143.74 | 11 | 1.99 | 0.064 | 0.166 |
|  | Vis.Easy | -4.31 ± 96.55 | 11 | -0.15 | 0.883 | 0.883 |
|  | Vis.Hard | -72.41 ± 151.51 | 11 | -1.77 | 0.116 | 0.186 |

Note. The table displays both permutation test-derived p-values and Benjamini-Hochberg FDR-adjusted q-values. Statistical significance thresholds were defined as follows: q < 0.05 (*) and q < 0.01 (**).

**Table S8 t- and p-values from paired T-tests comparing the TMS effect on ACC (%) between target site and correspondent control Vertex site at each subregion group (a-IFG, mid-IFG, a-pMTG, p-pMTG)**

|  | **Condition** | **ΔACC (M ± SD)** | **df** | **t** | ***p*_(Perm)_** | ***q*_(FDR)_** |
| --- | --- | --- | --- | --- | --- | --- |
| **a-IFG vs. Vertex** |  |  |  |  |  |  |
| (N = 7) | Sem.Easy | 1.43 ± 3.66 | 6 | 1.03 | 0.439 | 0.660 |
|  | Sem.Hard | 8.81 ± 9.16 | 6 | 2.54 | 0.030 | 0.181 |
|  | Vis.Easy | -0.48 ± 3.56 | 6 | -0.35 | 0.849 | 0.906 |
|  | Vis.Hard | -0.24 ± 4.95 | 6 | -0.13 | 1 | 1 |
| **mid-IFG vs. Vertex** |  |  |  |  |  |  |
| (N = 15) | Sem.Easy | 1.89 ± 3.20 | 14 | 2.28 | 0.031 | 0.181 |
|  | Sem.Hard | -1.33 ± 5.28 | 14 | -0.98 | 0.383 | 0.660 |
|  | Vis.Easy | 1.33 ± 4.04 | 14 | 1.28 | 0.174 | 0.464 |
|  | Vis.Hard | 2.11 ± 4.39 | 14 | 1.86 | 0.108 | 0.345 |
| **a-pMTG vs. Vertex** |  |  |  |  |  |  |
| (N = 10) | Sem.Easy | 0.33 ± 2.58 | 9 | 0.41 | 0.649 | 0.780 |
|  | Sem.Hard | 3.00 ± 9.36 | 9 | 1.01 | 0.388 | 0.660 |
|  | Vis.Easy | 0.17 ± 1.66 | 9 | 0.32 | 0.682 | 0.780 |
|  | Vis.Hard | 3.00 ± 3.41 | 9 | 5.44 | 0.034 | 0.181 |
| **p-pMTG vs. Vertex** |  |  |  |  |  |  |
| (N = 12) | Sem.Easy | 0.56 ± 3.04 | 11 | 0.63 | 0.523 | 0.698 |
|  | Sem.Hard | -2.08 ± 4.88 | 11 | 1-1.48 | 0.210 | 0.479 |
|  | Vis.Easy | 2.78 ± 4.68 | 11 | 2.06 | 0.060 | 0.241 |
|  | Vis.Hard | -1.53 ± 6.65 | 11 | -0.80 | 0.454 | 0.660 |

Note. Sixteen pairwise comparisons (paired t-tests and permutation test with Benjamini-Hochberg FDR correction) were conducted. The table displays both permutation test-derived p-values and Benjamini-Hochberg FDR-adjusted q-values. Statistical significance thresholds were defined as follows: q < 0.05 (*) and q < 0.01 (**).

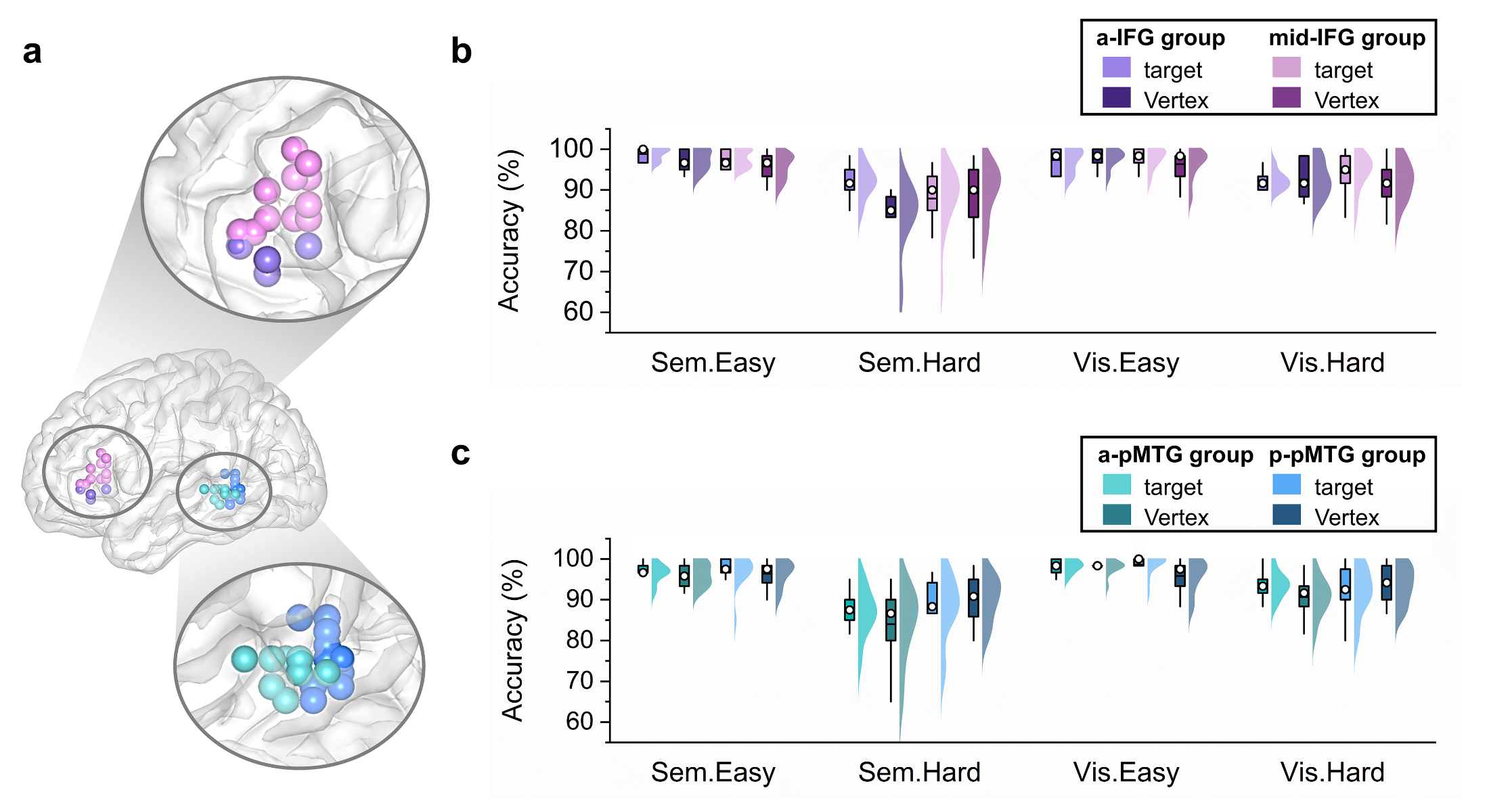

**Figure S1.** Behavioral differences in the TMS experiment between stimulation on the anterior and posterior subregions of the IFG and pMTG compared to Vertex stimulation. **(a)** Locations of TMS stimulation site are visualized on the standard ICBM152 template surface, with each site represented by a color-coded dot for each subject: aIFG in purple, mid-IFG in pink, a-pMTG in teal, and p-pMTG in blue. **(b)** Behavioral results for subregions of IFG under each task conditions, with participants divided into two groups based on stimulation site (a- and mid-IFG), and Vertex stimulation of the same group participants used as a comparison. **(c)** Behavioral results for subregions of pMTG under each task conditions, with participants divided into two groups based on stimulation site (a- and p-pMTG), with Vertex stimulation of the same group used as a comparison. The x-axis displays the four conditions, while the y-axis shows mean ACC of each condition. Boxplots show the median (denoted by hollow circles) and 1.5 times the interquartile range. Half-violin plots demonstrate the data distribution, with each region represented by a color matching the TMS stimulation site and corresponding Vertex represented by a darker color. Abbreviations: a-IFG, the anterior subregion of inferior frontal gyrus; mid-IFG, the middle subregion of inferior frontal gyrus; a-pMTG, the anterior subregion of posterior temporal gyrus; p-pMTG, the posterior subregion of posterior temporal gyrus; ACC, Accuracy. Statistical significance thresholds were defined as follows: *p* < 0.05 (*) and *p* < 0.01(**).

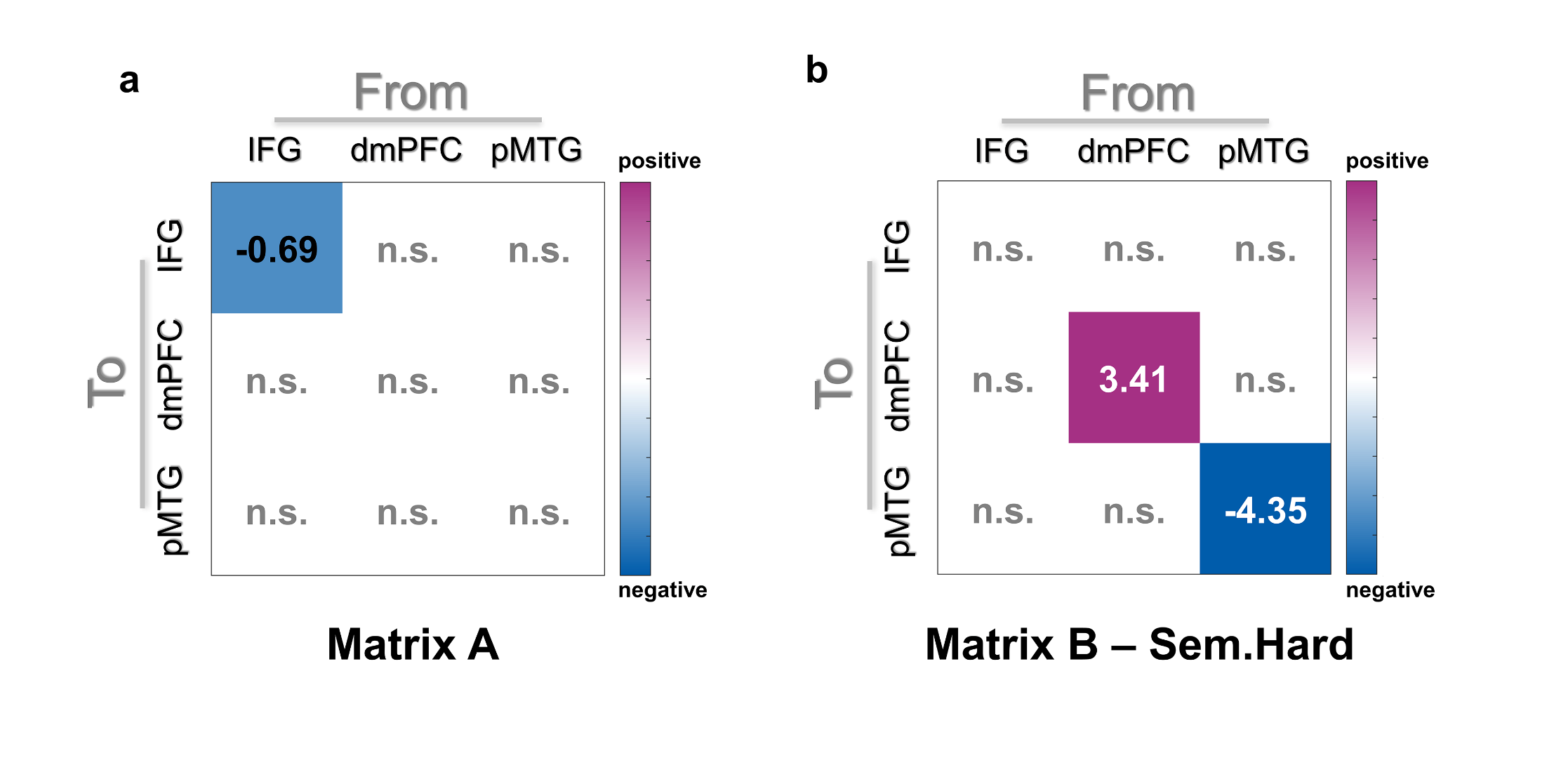

**Figure S2.** Effects of semantic control performance difference on connectivity parameters. **(a)** The matrix A: higher RV score (i.e., higher semantic control demand for the participant) was associated with the reduced autoinhibitory connection in the left IFG across all semantic conditions. **(b)** The matrix B: In the Sem.High condition, a higher RV score was associated with greater autoinhibition of the left dmPFC and further reduced autoinhibition of the left pMTG. Effective connectivity strengths are displayed by the color ranging from dark purple to white (indicating positive connectivity) and from white to dark blue (indicating negative connectivity). Parameters with stronger evidence (posterior probability > 95%) are presented and subthreshold parameters are marked with “n.s.”.
